# Selective bioelectronic sensing of quinone pharmaceuticals using extracellular electron transfer in *Lactiplantibacillus plantarum*

**DOI:** 10.1101/2023.03.23.533500

**Authors:** Siliang Li, Caroline De Groote Tavares, Joe G. Tolar, Caroline M. Ajo-Franklin

## Abstract

Redox-active small molecules containing quinone functional groups play important roles as pharmaceuticals, but can be toxic if overdosed. Despite the need for a fast and quantitative method to detect quinone and its derivatives, current sensing strategies are often slow and struggle to differentiate between structural analogs. Leveraging the discovery that microorganisms use certain quinones to perform extracellular electron transfer (EET), we investigated the use of *Lactiplantibacillus plantarum* as a whole-cell bioelectronic sensor to selectively sense quinone analogs. By tailoring the native EET pathway in *L. plantarum*, we enabled quantitative quinone sensing of 1,4-dihydroxy-2-naphthoic acid (DHNA) - a gut bifidogenic growth stimulator. We found that *L. plantarum* could respond to environmental DHNA within seconds, producing electronic signals that cover a 10^6^ concentration range. This sensing capacity was robust in different assay media and allowed for continuous monitoring of DHNA concentrations. In a simulated gut environment containing a mixed pool of quinone derivatives, this tailored EET pathway can selectively sense pharmacologically relevant quinone analogs, such as DHNA and menadione, amongst other structurally similar quinone derivatives. We also developed a multivariate model to describe the mechanism behind this selectivity and found a predictable correlation between quinone physiochemical properties and the corresponding electronic signals. Our work presents a new strategy to selectively sense redox-active molecules using whole-cell bioelectronic sensors and opens the possibility of using probiotic *L. plantarum* for bioelectronic applications in human health.

**Significant Statement:** Quinone-containing pharmaceuticals show toxicity at high concentrations, making it important to quickly and accurately measure their concentration while distinguishing between analogs. To address this problem, we leveraged recent discoveries in electroactive bacteria to develop a novel concept for whole-cell sensing. This concept combines selectivity and specificity, enabling differentiation between analogs based on the temporal dynamic of electron transfer in living cells. With this strategy, we achieved selective detection of pharmacologically relevant quinones with distinct electronic signals for each analog. These signals were deciphered by a multivariate model to provide insight into the specific physiochemical properties of each analog. We envision that this new concept can be applied to other analytes for faster and more efficient sensing using electroactive whole cells.

## Introduction

Redox-active small molecules facilitate interactions between different organisms and their environment. For example, redox-active molecules are tools for interspecies cooperation and competition (1), vehicles for organisms to gain nutrients and conserve energy from their environment (2), and messengers for engineering artificial cell-cell communication (3). A particularly significant set of redox-active small molecules contain the quinone functional group. These molecules, referred to as quinone, play a crucial role in biological energy transduction and have been widely used as pharmaceuticals to combat pathogens, tumors, and to reduce inflammation (4, 5). However, these quinone-derived drugs can cause adverse effects, including toxicity, at high doses or with long-term exposure (5, 6). Additionally, toxic naphthoquinone or benzoquinone derivatives released from industrial products or combustion of fossil fuels have contributed to air, water, and soil pollution, damaging the global ecosystem and posing a threat to human health (7, 8). Therefore, it is essential to accurately detect quinones and monitor their concentrations in order to properly utilize their beneficial properties while minimizing their toxic effects.

Existing strategies for detecting a specific quinone in biological samples typically involve high-performance liquid chromatography (HPLC) or liquid chromatography–mass spectrometry (LCMS) (9). However, these analytical methods use expensive equipment that requires highly-trained operators to perform calibration and sample preparation before any measurement. While electrochemical methods can also be used to detect and characterize quinones, they need to be coupled with analytical chromatography to differentiate quinone analogs (10). Thus, despite their importance, current technologies for selectively detecting a specific quinone in complex biological samples are slow, complex, and require equipment with high capital costs.

An emerging sensing technology that is fast, uses inexpensive instrumentation, and operates in complex samples is whole-cell bioelectronic sensors. These sensors can couple the presence of a chemical to an electronic response using microbes capable of transferring electrons to an electrode - a multi-step process known as extracellular electron transfer (EET) (11, 12). To create bioelectronic sensors, these microbes have been genetically manipulated to make specific steps in the EET path dependent on the presence of the target analyte using three strategies. The first strategy involves using transcriptional factors regulated by the analyte to conditionally express electron transfer proteins required for EET (13–15). While this approach is theoretically applicable to sensing various molecules, including quinones, the detection requires ex-situ analysis which slows the process to a minimum of 30 minutes. The second strategy selectively introduces or deletes an oxidoreductase that both consume the analyte as a substrate and affects the electron flux through the EET pathway. This strategy has been harnessed to sense electron donors (16), terminal electron acceptors (17), or oxyanions involved in assimilatory reduction (18). While this strategy can yield a real-time response, it cannot be applied to quinones since they are not consumed during EET. The third strategy involves engineering a protein in the EET pathway to be allosterically regulated by the analyte of interest. While this strategy has recently been used to sense an endocrine disruptor in real-time (18), it requires a known ligand-binding domain for the analyte of interest, which is not available for quinones. Thus, none of these strategies provide a rapid and selective way to sense quinones.

However, recent discoveries of the diverse ways EET is carried out in nature provide inspiration for new sensing strategies. Together with Light (19) and Marco (20), we recently discovered a novel EET pathway that is widely found in gram-positive Firmicutes, including a probiotic gut bacterium - *Lactiplantibacillus plantarum*. This lactic acid bacterium can reduce extracellular electron acceptors such as insoluble iron oxides and an extracellular electrode in the presence of exogenous quinone (20, 21). Here, we develop *L. plantarum* as a bioelectronic sensor for pharmacologically relevant quinones. We first examined its sensitivity, speed, and dose-dependence in response to DHNA (1,4-dihydroxy-2-naphthoic acid), a menaquinone precursor that can reduce gut inflammation and stimulate bifidogenic growth in the gut (22–24). We then expanded its ability to sense other quinone derivatives and found that *L. plantarum* can differentiate between different quinone analogs in a mixed quinone pool and generate distinguishable electronic signals. Finally, we tested the quinone bioelectronic sensor in a simulated gut fluid and explored the mechanism behind the quinone selectivity by creating a multivariate model. This work demonstrates a new strategy for rapidly and selectively sensing redox-active molecules and reveals the potential for using *L. plantarum* as a probiotic exoelectrogen for health-related applications.

## Results

### A tailored EET pathway in *L. plantarum* allows signal quantification

In order to develop *L. plantarum* as a bioelectronic sensor to sense quinone, it is necessary to ensure that the output electronic signal is directly proportional to the concentration of the quinone being sensed. We have shown previously that Ndh2, a membrane-bound type II NADH dehydrogenase, is required for EET in *L. plantarum* (20). The addition of exogenous quinones, such as DHNA, trigger Ndh2 to oxidize intracellular NADH and reduce exogenous quinones thereby directly or indirectly (via PplA and/or EetA) transferring electrons from the cytosol to the extracellular electron acceptors (20, 21) (Fig. 1A) . The sequential reaction can be described as below:

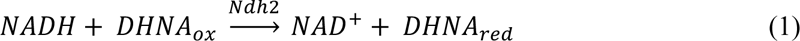

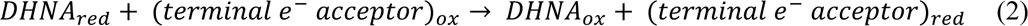

**Figure 1.**
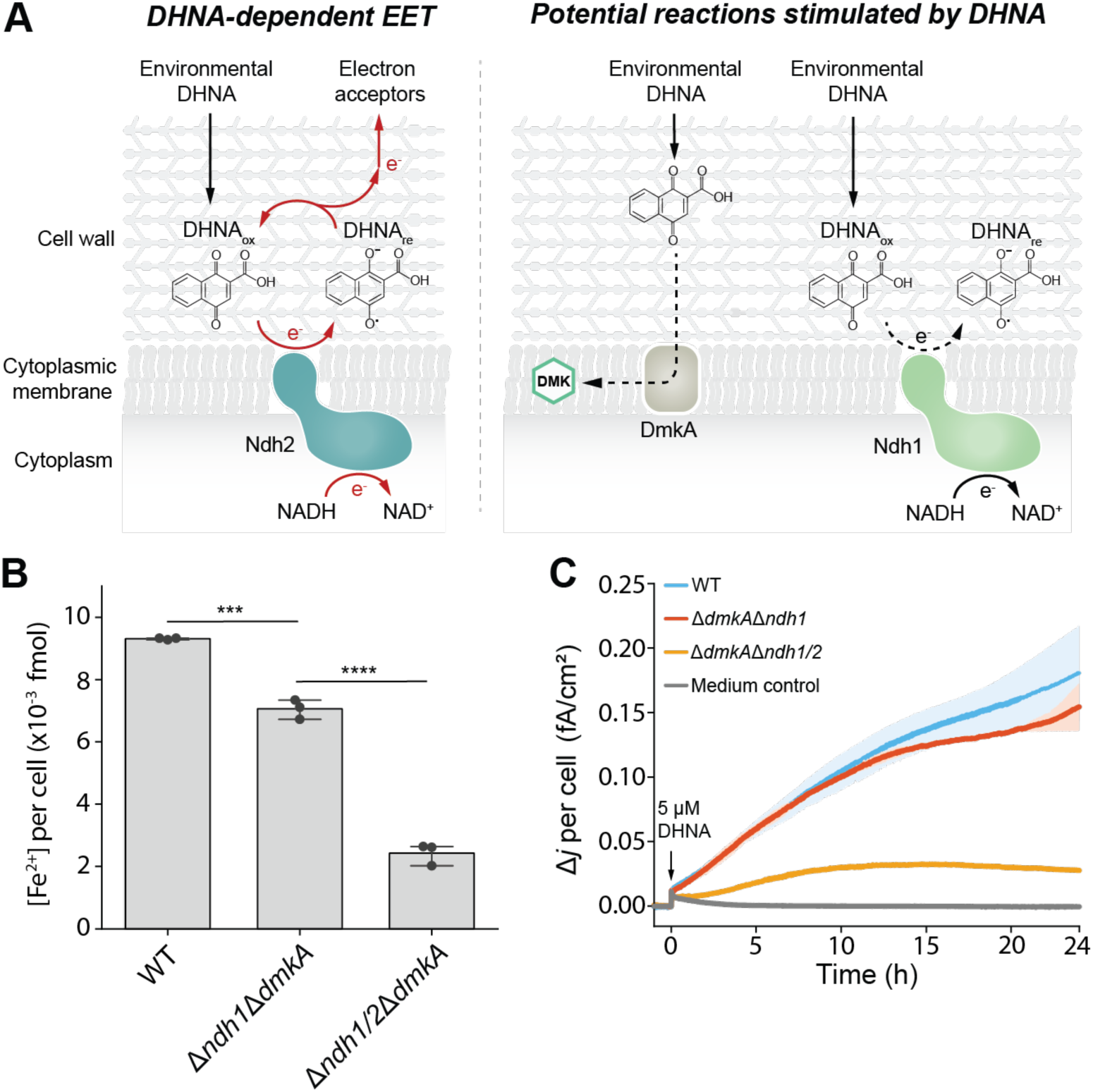
A tailored EET pathway in *L. plantarum* improves signal orthogonality in response to DHNA. (A) Schematic showing enzymatic reactions stimulated by environmental DHNA. Red arrows show that Ndh2-associated EET uses DHNA as electron shuttles. Dash arrows indicate the potential side reactions that occur when DHNA is present. (B-C) Comparing the EET activity of *L. plantarum* wide type (WT), side reactions removed mutant (Δ*dmkA*Δ*ndh1*), or Δ*dmkA*Δ*ndh1/2* mutant by either (B) iron reduction or (C) anode reduction. DHNA (5 μM) was added in all cases. For anode reduction, the electrode was polarized at +0.2 V v.s. Ag/AgCl. Δ*j* per cell represents the delta current density normalized to CFU. Δ*j* of medium control was also divided by the mean CFU to bring values to the same magnitude for comparison. Data represent mean ± 1 s.d. for n=3 (B) or n=2 (C) biological replicates. Significant differences were determined by one-way ANOVA with Tukey’s test, ***p ≤ 0.001, ****p ≤ 0.0001.

We hypothesized that, at steady state, the overall EET rate is determined by reaction 1 and follows Michaelis-Menten kinetics (equations 3-5 in Materials and Methods). Therefore, the DHNA concentration is directly linked to EET, demonstrating the possibility to use *L. plantarum* to quantitatively sense quinone. To experimentally test this working model, it is required that the observed electronic signal is solely driven by Ndh2. However, enzymes that are not essential for EET, including DmkA, an octaprenyltransferase that converts DHNA to dimethylmenaquinone (DMK) (19), and Ndh1, a type II NADH dehydrogenase that has a role in the aerobic-respiration-like response of *L. plantarum* (25), also speculated to react with DHNA. These reactions could potentially interfere with DHNA reduction driven by Ndh2 (Fig. 1A).

We sought to determine if this interference could be eliminated by generating a tailored EET pathway. To do so, we deleted *dmkA* and *ndh1* from *L. plantarum* NCIMB8826. A mutant with an additional *ndh2* deletion served as the control to confirm the role that Ndh2 plays in EET. We compared the EET activity of the mutants by testing their ability to reduce either iron(III) oxide (Fe^3+^ to Fe^2+^) or anode (produce current) in the presence of 5 μM DHNA. The Δ*dmkA*Δ*ndh1* strain showed slightly decreased levels of iron reduction and current generation compared with the wild-type strain, likely because of branched or secondary EET pathways associated with DmkA or Ndh1. The additional *ndh2* knockout significantly disrupted the EET activity (Fig. 1B-C). These results are consistent with our hypothesis that the majority of the electronic signal is driven by Ndh2 whereas DmkA and Ndh1 are dispensable for EET activity. Thus, we used the Δ*dmkA*Δ*ndh1* mutant strain in the following experiments to test how *L. plantarum* responds to environmental quinones.

### *L. plantarum* sensitively responds to a wide range of DHNA concentrations following Michaelis-Menten kinetics

To establish the potential of *L. plantarum* to serve as bioelectronic sensors for quinones, we sought to first determine the sensitivity, dynamics, and cell-dependence of the EET in response to physiologicolly-relevant concentrations of DHNA. Therapeutically, 5 nM DHNA produced by food-derived propionibacteria can stimulate the growth of bifidobacteria in vitro (22), 0.1 μM DHNA exhibits anti-inflammatory activity in vitro (23), and 2 mg/kg DHNA inhibits colitis in mice model (23). However, a high concentration of DHNA (50-100 μM) induces significant cytotoxicity in mouse and human cell lines (24). Under well-controlled laboratory conditions, microbes can synthesize DHNA to even higher concentrations, ranging from 1.8 - 236 μM (26–28). To determine how sensitively *L. plantarum* responds to environmental DHNA, we incubated the Δ*dmkA*Δ*ndh1* strain with varied DHNA concentrations and measured iron reduction levels at several time points over 12 hours. We observed that *L. plantarum* could react with DHNA over a 10^6^ concentration range from 5 nM to 500 μM (Fig. 2A, 2B), indicating a highly sensitive EET machinery with a large dynamic range. To analyze if EET can quantitatively reflect DHNA concentration following Michaelis-Menten kinetics, we calculated the iron reduction rate per cell over 12 h for each DHNA concentration (Fig. S1). The data could be fitted into the Michaelis-Menten equation (Fig. 2C) (equations 4-5 in the Materials and Methods), which confirms our working model. Moreover, the iron reduction rate remained constant over the experimental period, as can be seen from the linear increase in iron reduction levels (Fig. S1), implying that the reaction did not reach equilibrium over 12 h. These data suggest that *L. plantarum* can serve as a quantitative bioelectronic sensor for DHNA.

**Figure 2.**
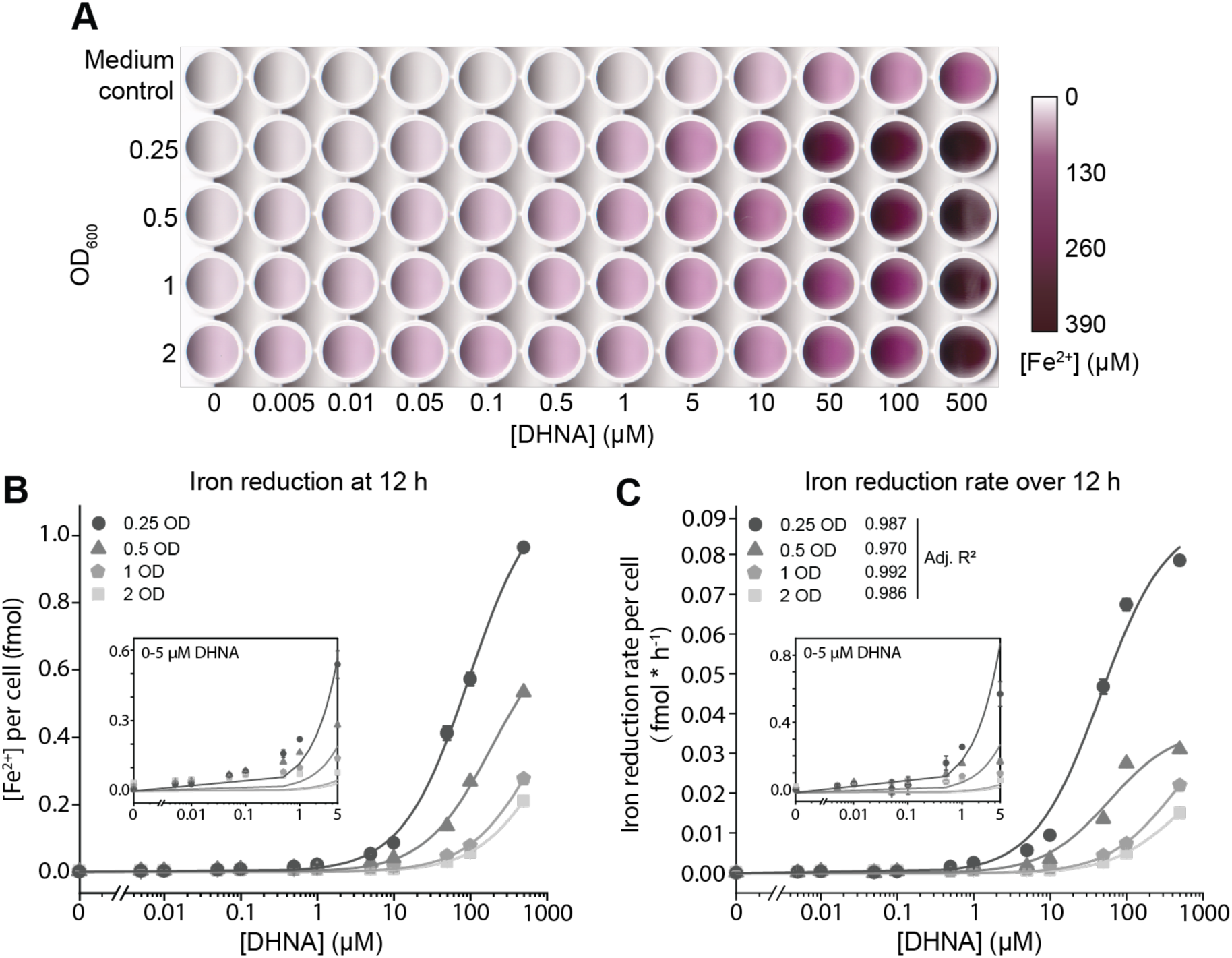
EET of *L. plantarum* shows high sensitivity and dynamic range when responding to environmental DHNA. (A) Scanned image showing iron reduction levels when different density of cells was incubated with 0-500 μM DHNA for 12 h. The purple color represents Fe^2+^ concentration in each well. Medium control indicates DHNA-induced abiotic iron reduction. (B) Quantitative analysis of iron reduction levels in (A). Biotic iron reduction of each well was calculated by subtracting abiotically produced Fe^2+^ from biotically produced Fe^2+^ and normalizing to CFU of the inoculated cell. Lines are guides to the eye. The inset shows an enlarged region for 0-5 μM DHNA. Data represent mean ± 1 s.d. for n=3 biological replicates. (C) Iron reduction rate per cell (i.e. initial velocity of iron reduction from 0 h to 12 h normalized to CFU, see also Fig S1) as a function of DHNA concentration. Data were fitted into the Michaelis-Menten equation. The inset shows an enlarged region for 0-5 μM DHNA.

The Michaelis-Menten equation also implies a correlation between Ndh2 concentration and EET rate (equations 4-5 in the Materials and Methods). Since the total amount of Ndh2 in a given culture volume should depend on the number of cells, we sought to explore how cell density affects iron reduction and to determine an optimized cell density that is suitable for sensing. We found that, for all the tested DHNA concentrations (0-500 μM), the iron reduction level at 12 h and the iron reduction rate over 12 h both decreased when the optical density (OD) of cells increased (Fig. 2B, C). This effect could be attributed to the decreased carbon source and electron acceptor available for a single cell caused by the increased cell density in a fixed culture volume, which modulates EET rate at the single cell level. Since OD = 0.25 showed the largest fold change in EET in response to DHNA (Fig. 2B, C), this fixed cell density was chosen for quinone sensing.

### Temporal dynamics of EET reflect the concentration of environmental DHNA in real-time

Bioelectronic sensors can be created by integrating EET capable cells into bioelectrochemical systems (BESs) - hybrid biological and electrochemical systems that allow continuous monitoring of electronic signals. The response speed of bioelectronic sensors can vary substantially, from minutes (18) to hours (13, 14), depending on the sensing strategy. To investigate how rapidly *L. plantarum* responds to environmental quinones, we inoculated the Δ*dmkA*Δ*ndh1* strain at the OD = 0.25 into the BESs and monitored the temporal dynamics of the electronic signal when physiological concentrations of DHNA (0-5 μM) were injected. We probed the electronic signal by calculating the difference in current density produced on a per cell basis before and after DHNA injection (Δ*j* per cell). As a first test, the nutrient-rich chemically defined media (CDM) was used as the medium to support growth. Immediately upon the injection of DHNA into BESs, we observed a transient current spike (Fig. 3A). This transient spike was due to the abiotic oxidation of DHNA by the electrode, as a similar spike caused by DHNA was also observed in our previous study when heat-killed cells were inoculated (21). Following the spike, the DHNA triggered EET within seconds, and a higher DHNA concentration resulted in a higher current magnitude (Fig. 3A, left). The biotic current produced by EET then gradually increased over 24 hours (Fig. 3A, right). We found that the current density per *L. plantarum* cell could differentiate between DHNA concentrations with 99% confidence (*p* < 0.01) within 21 seconds, highlighting the real-time response of the DHNA-induced EET (Fig. 3B). To further dissect the temporal dynamics, we used linear regression to calculate the initial rate of current increase within 2 hours (Fig. S2A). The increasing rate is also correlated with DHNA concentration (Fig. S2B). Together, these data suggest that DHNA can dose-dependently activate EET, and the temporal dynamics of current produced by EET reflect DHNA concentrations in real time.

**Figure 3.**
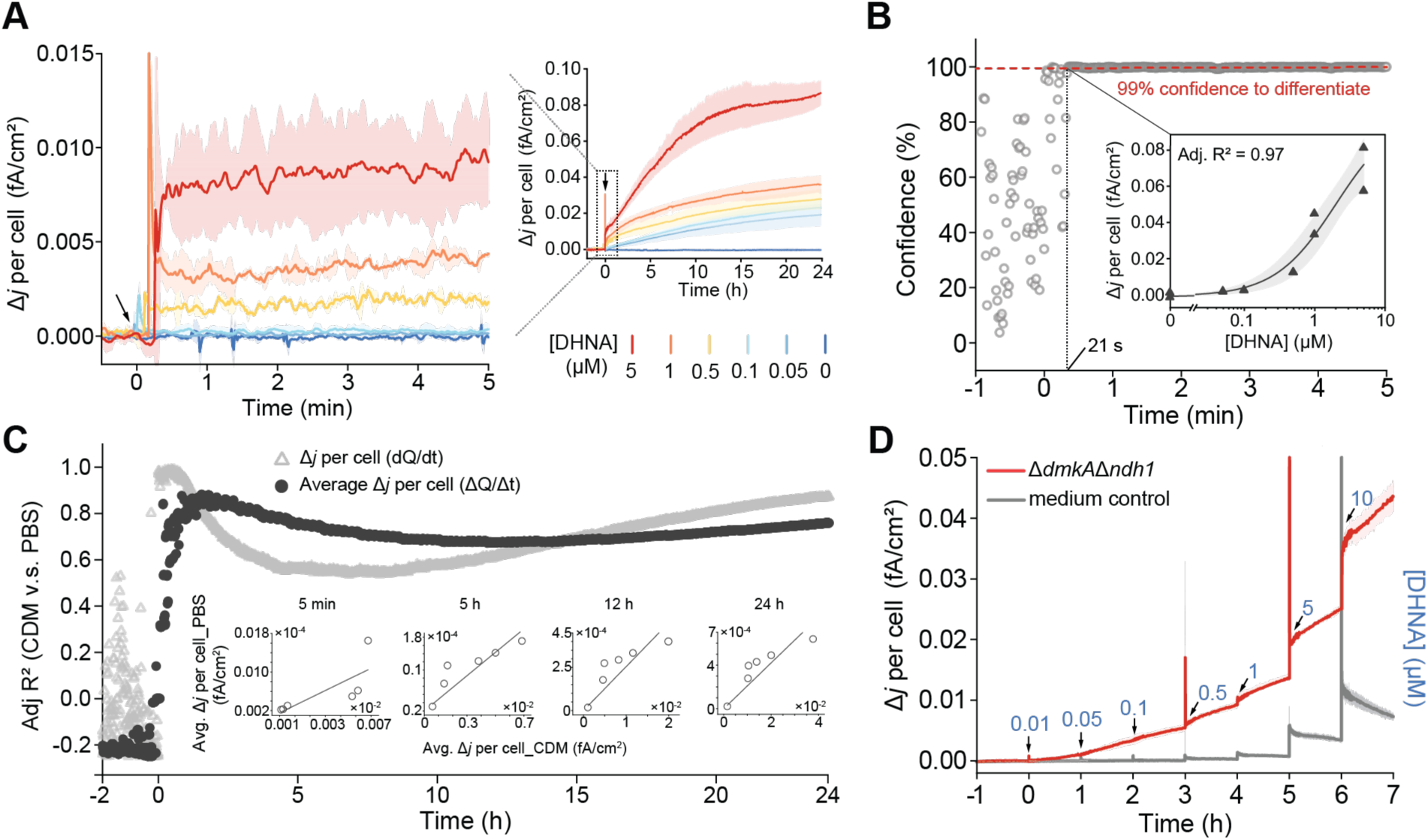
*L. plantarum* rapidly responds to DHNA with dose-dependent temporal dynamics. (A) Current density produced by *L. plantarum* Δ*dmkA*Δ*ndh1* strain upon exposure to different concentrations of DHNA in bioelectrochemical systems (BESs) within 24 h (right panel). An enlarged region showing rapid current response within 5 minutes (left panel). The black arrow indicates DHNA injection. Data represent mean ± 1 s.d. for n=2 biological replicates. (B) Confidence to differentiate varied DHNA concentrations as a function of time. Confidence was determined by one-way ANOVA for current densities at each single time point. The data in the inset were fitted into the Michaelis-Menten equation. (C) Comparison of dose-dependent temporal dynamics in response to DHNA when tested in CDM or PBS. The hollow triangles represent the comparison of absolute level of current density (dQ/dt per cm^2^). The black dots represent the comparison of average current density (ΔQ/Δt per cm2). Insets showcase the results of comparison of average current density at 5 min, 5 h, 12 h, and 24 h. Each dot in the insets represents a different DHNA concentration. The mean data of n=2 biological replicates for both CDM and PBS were used for comparison. (D) Current density produced by *L. plantarum* upon exposure to stepwise increasing concentrations of DHNA. Medium control indicates the abiotic current upon DHNA injection. Data represent mean ± 1 s.d. for n=2 biological replicates.

To test the EET dynamics when cells were in a resting, rather than growing state, we repeated the above experiment in phosphate buffered saline (PBS), which lacks nutrients for protein synthesis and cell proliferation. Again, EET was dose-dependently activated by DHNA within seconds after DHNA injection (Fig. S3A-B). The varied DHNA concentrations could be differentiated with 99% confidence within 36 seconds (Fig. S3C). Interestingly, the biotic current reached a significantly smaller plateau (∼5×10^-4^ *vs*. ∼2×10^-2^ fA/cm^2^ per cell) in less time (∼7.5 h vs. ∼24 h) under these resting conditions, suggesting that cell proliferation and/or gene expression increase EET. Nonetheless, these results demonstrate that *L. plantarum* can sense and respond to DHNA in a dose-dependent manner in both nutrient-rich and nutrient-poor media.

Effective bioelectronic sensors accurately report the concentration of analyte independent of other physicochemical variations in the sample. To rigorously investigate the reliability of DHNA sensing in different media, we compared the current produced by the *L. plantarum* Δ*dmkA*Δ*ndh1* strain in PBS or CDM in response to different concentrations of DHNA, and used adjusted R^2^ to determine the data similarity. In the course of 24 h post DHNA injection, we found a consistent moderate to high adjusted R^2^ between these two datasets (Fig. S4), suggesting that the temporal dynamics of current produced by EET could reproducibly reflect DHNA concentrations irrespective of media components. However, the adjusted R^2^ started to fluctuate over 24 h after reaching the plateau within 5 min (Fig. S4). This suggests that, although data from different media can be aligned, the absolute level of current density (dQ/dt per cm^2^) for a fixed DHNA concentration is still dependent on whether the cells are in PBS or CDM. As an alternative, we calculated the average current density (ΔQ/Δt per cm^2^) to focus on the temporal evolution of current change rather than the absolute current level. We observed an improved consistency between the two datasets by using this analysis (Fig. 3C). We conclude that, although the absolute level of current density is suitable for short-term sensing, the average current density is more suitable for long-term monitoring of DHNA. This provides a methodology for data comparison in bioelectronic sensing.

We next sought to explore if *L. plantarum* could continuously sense environmental quinone by adding incremental concentrations of DHNA in a stepwise manner with CDM as the assay medium. With every successive addition of a higher concentration of DHNA at a one-hour interval, we observed a steeper current density in response (Fig 3D). The speed of the current increase correlates with the accumulated DHNA concentrations (Fig. S2C). Again, there is a transient current spike induced by DHNA injection starting at 1 μM DHNA (Fig 3D). The spike magnitude showed no difference between EET and medium control, confirming this signal arises from the abiotic reaction between DHNA and electrode (Fig. S5). Following the spike, the current for medium control gradually went down, indicating the increased current for the Δ*dmkA*Δ*ndh1* strain was solely driven by EET (Fig 3D). These results suggest that the *L. plantarum* can continuously sense DHNA, adjust its EET activity accordingly, and report the DHNA concentration as electronic signals in real-time.

### *L. plantarum* selectively senses pharmacological-relevant quinones in a complex environment

In the preceding experiments, we have reported on the sensitivity, rapidity, dynamics, reproducibility, and continuity of DHNA sensing by *L. plantarum*. However, quinones encompass a large group of redox-active aromatic compounds derived from core structural motifs, with DHNA belonging to the class of 1,4-naphthoquinone (1,4-NQ) derivatives. It has been reported that *L. plantarum* can utilize different 1,4-NQ derivatives to support EET (29). However, the selectivity among these derivatives remains unknown. Therefore, we next investigated if *L. plantarum* could discriminate specific quinone derivatives from an environmental quinone pool containing diverse structural analogs, and thus, behave as a selective quinone bioelectronic sensor (Fig. 4A). We tested six quinone derivatives (one 1,4-benzoquinone and five 1,4-NQ derivatives) that have pharmacological effects. These derivatives were hydroquinone (depigmenting agent) (30), 1,4-NQ (anti-cancer agent) (31), DHNA (anti-inflammation agent, bifidogenic growth stimulator) (22–24), menadione (vitamin K precursor) (32), juglone (herbicide) (33), and DCDMQ (anti-cancer agent) (34). With the assumption that Ndh2 may play a role in quinone recognition, we injected the six quinone derivatives individually to either the Δ*dmkA*Δ*ndh1* or the Δ*dmkA*Δ*ndh1/2* strains and monitored the differential current density produced by these two mutants. Intriguingly, the Δ*dmkA*Δ*ndh1* strain produced more current when reacting with DHNA or menadione compared to the Δ*dmkA*Δ*ndh1/2* strain, while no difference was observed when reacting with other quinone derivatives (Fig. S6A). This observation indicates that Ndh2 can specifically recognize DHNA and menadione to support EET.

**Figure 4.**
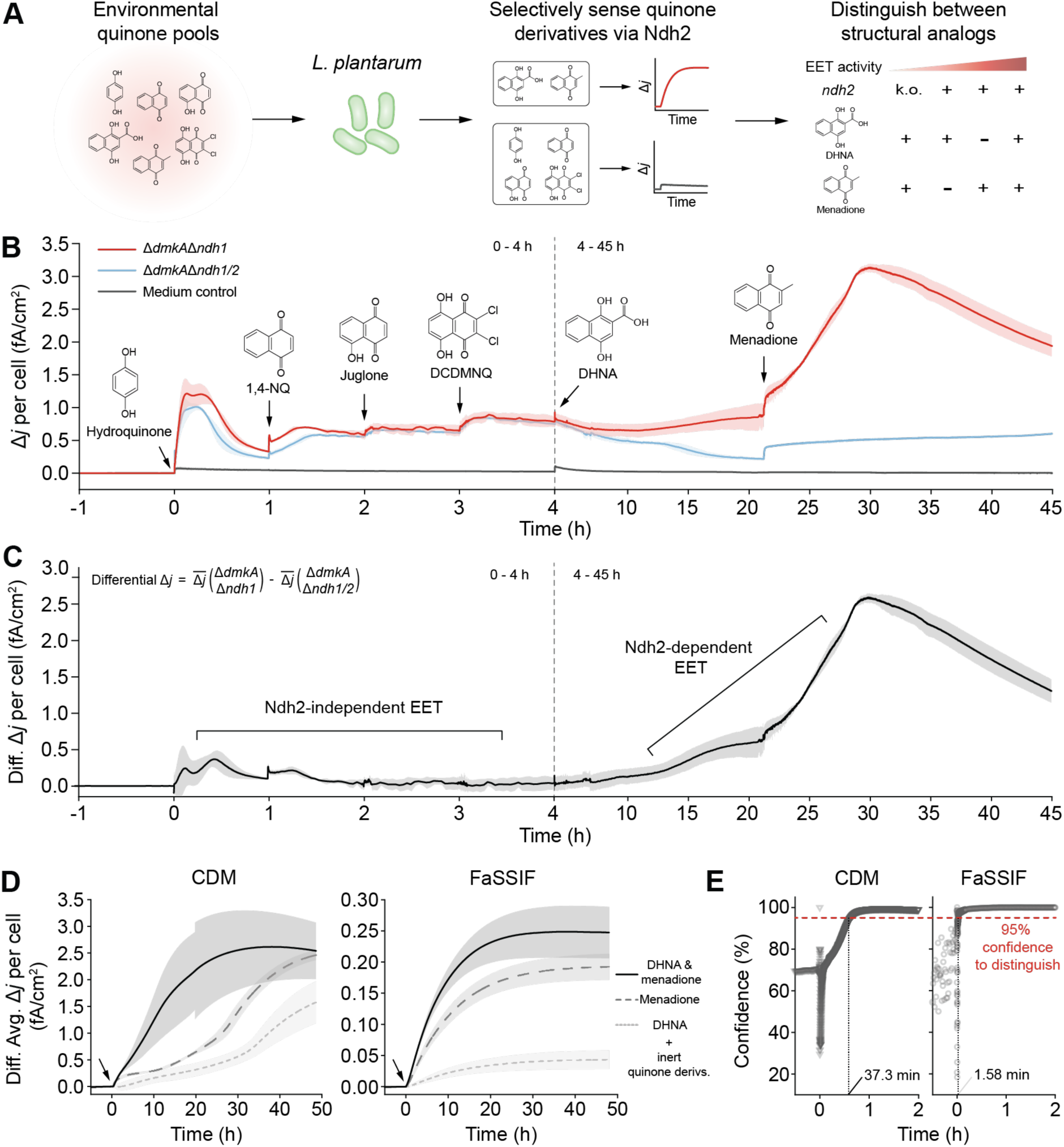
Ndh2-dependent EET discriminates quinone structural analogs. (A) Illustration showing Ndh2-dependent EET of *L. plantarum* selectively senses specific quinone derivatives from an environmental quinone pool. (B) Current density produced by *L. plantarum* Δ*dmkA*Δ*ndh1* or Δ*dmkA*Δ*ndh1/2* mutant compared to medium control when 1 μM of hydroquinone or 1,4-NQ derivatives were added stepwise. The next compound was not injected until the current of the prior compound stabilized. Data represent mean ± 1 s.d. for n=2 biological replicates. (C) The differential current density was calculated by subtracting the mean Δ*j* per cell of Δ*dmkA*Δ*ndh1/2* strain from that of Δ*dmkA*Δ*ndh1* strain. The shaded error bar represents propagated error. (D) Differential average current density produced by *L. plantarum* when 1 μM of DHNA, menadione, or both were added to the background of inert quinone derivatives (hydroquinone, 1,4-NQ, juglone, DCDMNQ). Inert quinone derivatives were added at 1 μM in CDM and 0.01 μM in FaSSIF. Data represent mean ± propagated error for n=2 biological replicates. (E) Confidence to distinguish between DHNA, menadione, or both as a function of time. Confidence was determined by one-way ANOVA on data shown in (C).

To test if *L. plantarum* can selectively sense DHNA and menadione in a continuous way, we injected the six quinone derivatives sequentially into the same batch of cells and monitored the temporal dynamics of current density. When hydroquinone, 1,4-NQ, juglone, or DCDMNQ were sequentially added, the two mutants produced a similar current in response (Fig. 4B). However, when DHNA and menadione were further injected, the Δ*dmkA*Δ*ndh1* strain produced more current than the Δ*dmkA*Δ*ndh1/2* strain (Fig. 4B). To isolate the signal for Ndh2-dependent EET, we calculated the differential current density by subtracting the current produced by the Δ*dmkA*Δ*ndh1/2* strain from that produced by the Δ*dmkA*Δ*ndh1* strain (Fig. 4C). The Ndh2-independent EET does not significantly affect the differential current density, while the Ndh2-dependent EET results in an increased differential current density. Moreover, the injection of menadione resulted in a more rapid increase in the differential current density than DHNA. In this way, DHNA and menadione could be further differentiated (Fig. 4C). This result also implies that sensing the first quinone did not attenuate the ability to sense the second one. Thus, although several quinone structural analogs were present in the same environment, the Ndh2-dependent EET in *L. plantarum* can continuously and selectively sense EET-supportive quinone derivatives and produce distinguishable signals.

We further explored if *L. plantarum* could distinguish DHNA and menadione from the other quinones when they were all injected together. In addition to CDM, we also tested in a fasted state simulated intestinal fluid (FaSSIF) to explore *L. plantarum*’s sensing ability in a simulated gut environment (35). The inert quinone derivatives (hydroquinone, juglone, DCDMNQ, 1,4-NQ) were added at a lower concentration (0.1 μM) in FaSSIF than in CDM (1 μM) to mimic the physiological quinone background in the human gut. To these background media, we then added DHNA, menadione, or both and calculated the differential average current density to neutralize signal fluctuations and analyze the Ndh2-dependent EET (Fig. S7). In both media, we observed the highest differential average current density when DHNA and menadione were added together, followed by menadione and DHNA alone (Fig. 4D). This data demonstrates that DHNA and menadione have an additive effect in activating Ndh2-dependent EET. Thus, *L. plantarum* can produce distinguishable electronic signals in response to different quinone combinations in a mixed quinone pool. The time needed to differentiate DHNA, menadione, and the mixture with 95% confidence was 37.3 min and 1.58 min in CDM and FaSSIF, respectively (Fig. 4E). The longer time needed in CDM is attributed to the transient Ndh2-independent EET induced by the inert quinone derivatives that masked the signals from Ndh2-dependent EET (Fig. S7). In an environment with lower background, such as in FaSSIF, the time needed to differentiate DHNA and menadione is ∼20-fold lower.

### Mechanism of *L. plantarum* as a selective quinone bioelectronic sensor

Based on all our results, we sought to dissect the fundamental principles of how *L. plantarum* behaves as a selective bioelectronic sensor. We treated the reaction between quinone and the Ndh2-dependent EET pathway as a series of four successive steps: 1) quinone entering the cell membrane, 2) binding to Ndh2, 3) transferring electrons from the co-factor flavin adenine dinucleotide (FAD), 4) semi or hydroquinone leaving Ndh2 and transferring electrons through the rest of the EET pathway (Fig. 5A). We hypothesized that quinones could be selected by *L. plantarum* for EET through these four steps based on their physiochemical properties (Fig. 5B). These properties include lipophilicity/hydrophilicity, which determines if quinone can freely move in and out of the cell membrane, binding affinity, which determines how tight quinone binds to Ndh2, and redox potential, which determines the tendency to acquire and lose electrons.

**Figure 5.**
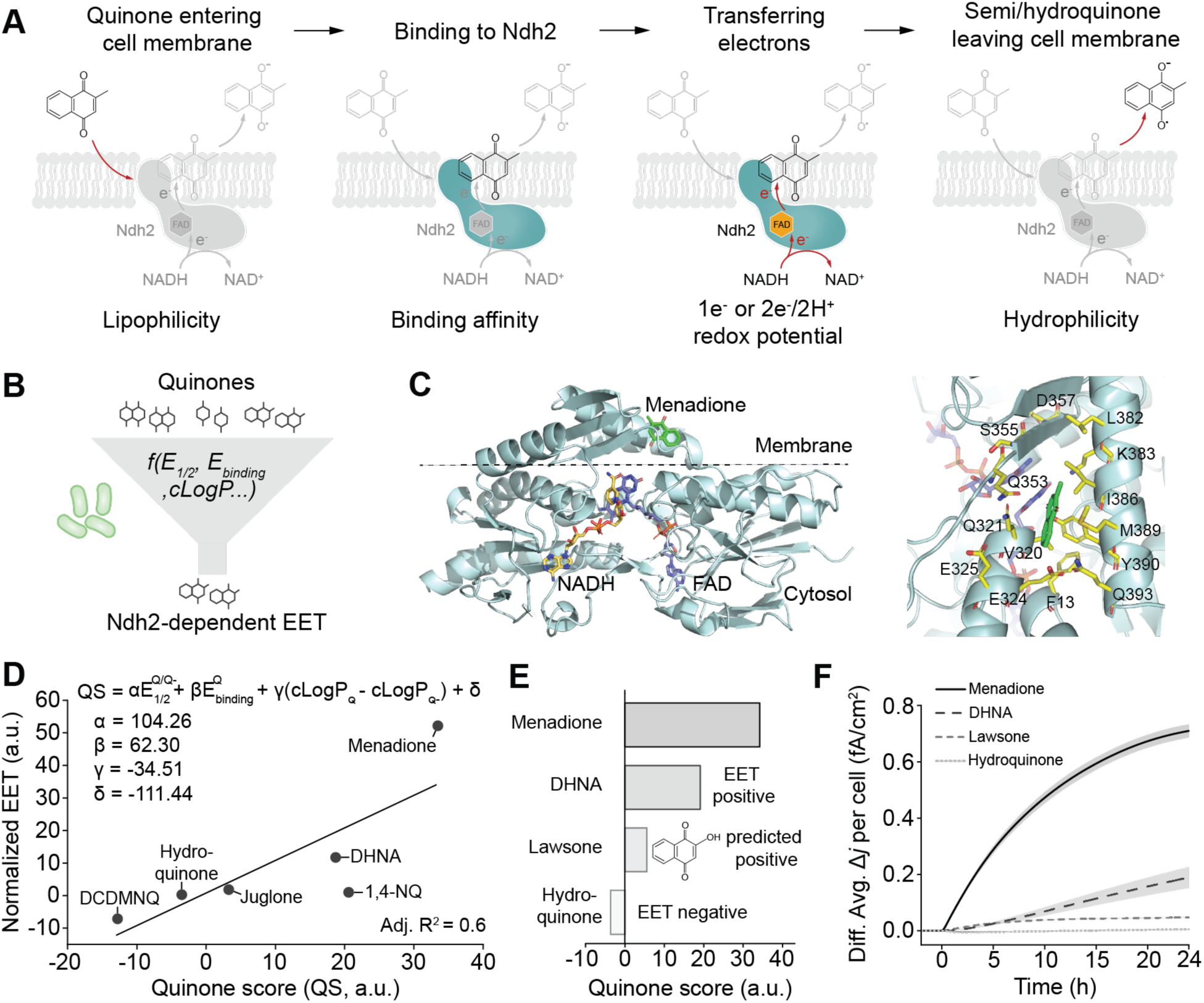
Mechanism of *L. plantarum* as a selective quinone bioelectronic sensor. (A) A four-steps model denoting the reaction between quinone and Ndh2 of *L. plantarum*. Below each step shows the potential properties of a quinone that would determine whether this quinone will be selected for Ndh2-dependent EET. (B) *L. plantarum* as a bioelectronic sensor to sense environmental quinones, process physiochemical information, and selectively output electronic signals in response to quinones that favor EET. (C) Predicted *L. plantarum* Ndh2_1-409_ structure, simulated quinone-Ndh2 binding, and the quinone binding pocket. (D) Correlation between normalized EET and the quinone score. 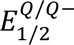, quinone 1 e^-^ redox potential; 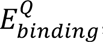, predicted binding energy between quinone and Ndh2; (𝑐𝐿𝑜𝑔𝑃_’_ -𝑐𝐿𝑜𝑔𝑃_’(_), differential of the calculated partition coefficient between quinone and semiquinone. (E) Lawsone (2-hydroxy-1,4- naphthoquinone) has a predicted quinone score greater than the negative control but smaller than DHNA. We predicted lawsone to be EET positive. (F) Differential average current density generated by *L. plantarum* when reacting with 1 μM menadione, DHNA, lawsone, or hydroquinone. Data represent mean ± propagated error for n=2 biological replicates.

To quantitatively analyze these properties, we determined the following three parameters for each quinone derivative: the calculated partition coefficient (cLogP) (Materials and Methods) (36); the predicted binding free energy between quinone and Ndh2 (Materials and Methods, Fig. 5C, S8) (37); and either the experimentally measured or predicted redox potential (Materials and Methods, Fig. S9, S10). All parameters are shown in Table S1. We then plotted these parameters against the differential average current density induced by each quinone derivative (based on the data shown in Fig. S6A) to analyze the correlations between EET and quinone parameters (Fig. S11). For redox potential, previous work shows that quinones prefer to undergo one electron transfer (1e^-^) in an aprotic environment (quinone⇋semiquinone, Fig. S9C), such as in the cell membrane, and revealed a correlation between quinone 1e^-^ redox potential and biophotocurrent generated by purple bacteria (38). Indeed, we found a weak correlation between Ndh2-dependent EET and 1e^-^ redox potential (adj R^2^ = 0.33) (Fig. S11A), while no correlation was observed for 2H^+^/2e^-^ redox potential (adj R^2^ = 0.13) (Fig. S11B). However, 1e^-^ redox potential alone fails to explain why juglone, 1,4-NQ, and hydroquinone, which have a lower 1e^-^ redox potential than DHNA, cannot be utilized by EET. For quinone-Ndh2 interaction, we first used AlphaFold to predict the Ndh2 structure for *L. plantarum* (Fig. S8) (39). Then, we used AutoDock Vina to simulate quinone-Ndh2 binding (37). We were able to predict a similar quinone binding pocket in the Ndh2 structure of *L. plantarum* compared to the homologs in other bacteria or yeast (40, 41), demonstrating the reliability of our simulation methods (Fig. 5C). However, no obvious correlation was found between the binding energy and EET (adj R^2^ = −0.23) (Fig. S11C). For lipophilicity/hydrophilicity, the cLogP value of quinone or semiquinone also fails to correlate with EET (adj R^2^ = −0.23, −0.24, respectively) (Fig. S11D, E). Thus, we concluded that a single parameter is insufficient to describe the relationship between quinone and Ndh2-dependent EET. This finding is not wholly surprising given that our mechanistic model suggests all of these three parameters would affect EET.

Inspired by previous work (38, 42, 43), we performed a multivariate analysis by integrating all parameters. To decrease the variate number, we calculated the differential of cLogP between quinone and semiquinone and treated it as a single variate (cLogP_Q_-cLogP_Q-_) (Fig. S12F). We first normalized the current density and all the variates to the unsubstituted 1,4-NQ (i.e., all variates for 1,4-NQ=1). Then, multivariate regression was performed and we termed the model output as “quinone score”. We observed a positive correlation between the quinone score and the normalized EET (adj R^2^ = 0.6) (Fig. 5D), suggesting that the multivariate model is able to interpret the quinone-EET relationship. However, some of the outliers cannot be explained by the model, suggesting that there are still parameters that our model does not cover. For example, 1,4-NQ has a relatively high quinone score but cannot be utilized by Ndh2-dependent EET.

Nonetheless, to test the predictive power of our model, we predicted the quinone score for lawsone (2-hydroxy-1,4-naphthoquinone) - a natural hair dye present in henna plants with a relatively low 1e^-^ redox potential (44, 45). The quinone score for lawsone is higher when compared to the EET- negative control but lower than those for DHNA and menadione (Fig. 5E). This score predicts that *L. plantarum* should be able to use lawsone as an electron shuttle in its Ndh2-dependent EET pathway, but will generate a lower current compared to DHNA or menadione. Indeed, when lawsone was supplied, the Δ*dmkA*Δ*ndh1* strain produced more current than the Δ*dmkA*Δ*ndh1/2* strain (Fig. S6B). When comparing the differential average current densities, the magnitude is higher for lawsone than that for hydroquinone (negative control) but lower than those for DHNA or menadione (Fig. 5F), which is in agreement with the prediction. Thus, our multivariate model could identify new quinone derivatives that can be sensed by Ndh2-dependent EET. Using this model, the electronic signals produced by *L. plantarum* can be decoded to distinguish between quinone structural analogs.

## Discussion

Here we investigated the potential of *L. plantarum* to serve as a bioelectronic sensor for quinones. Using a genome-modified *L. plantarum* strain, we demonstrated its ability to sensitively, rapidly, and continuously sense varying concentrations of DHNA in both nutrient-rich and nutrient-poor media following Michaelis–Menten kinetics. Further analysis revealed that the Ndh2-dependent EET in *L. plantarum* could selectively sense menadione, DHNA, and lawsone amongst other quinone structural analogs, and we created a multivariate model to explain this selectivity.

Our multivariate model reveals general principles underlying *L. plantarum*’s ability to selectively sense quinones. The model suggests that Ndh2-dependent EET mainly discriminates based on 1e^-^ redox potential of quinones, followed by its binding affinity to Ndh2, and least on its lipophilicity or hydrophilicity (Fig 5D). The positive and negative coefficients of the multivariate equation also imply that, in general, quinones with a higher tendency to lose electrons (lower 1e^-^ redox potential), higher binding affinity to Ndh2, and a smaller differential in lipophilicity/hydrophilicity (so that can easily move in and out the cell membrane) are more likely to be utilized by *L. plantarum* for EET. Compared to other models that examine the structure-function relationships between mediator and EET (38, 43), our model considers the biological interaction between quinone and a specific reductase (Ndh2) in addition to the quinone’s physiochemical properties. However, our model is limited by being based on a relatively small set of 1,4-naphthoquinones with similar physio-chemical properties. Thus, while this model has a degree of predictive power, future work is needed to refine and validate the model with a larger pool of quinone derivatives.

Our study reveals a new whole-cell bioelectronic sensor chassis - *L. plantarum*, and a strategy to sense a new class of analytes - quinones. State-of-the-art whole-cell bioelectronic sensors rely on *Shewanella oneidensis* (13–15), *Geobacter sulfurreducens* (17), or lab-engineered *Escherichia coli* (18), and are limited in their applications to environmental sensing. By demonstrating that *L. plantarum*, a probiotic gut microbe, can sense quinones in a simulated intestinal fluid (Fig. 4C), our study expands the potential applications of bioelectronic sensors to include sensing within mammalian hosts. Moreover, we characterized quinones as a new class of redox analytes that can be sensed by exoelectrogens to produce amperometric signals. Several prior studies have used redox-active molecules, such as flavins (46, 47) and phenazines (48), as signal carriers. However, most of them rely on voltammetric approaches where the signal generated by other redox molecules present in the medium often overlaps with the signal generated by the analytes. In contrast, *L. plantarum*’s sensing of quinone through a biologically selective EET route allows for more accurate sensing. Since quinone is recycled by EET, there is no change in the environment during the process of sensing. This is in contrast to other EET-based bioelectronic sensors that consume donors/acceptors or oxyanions during the sensing process (16–18). The recycling of quinone also amplifies the signals, making the limited detection concentration of our bioelectronic sensor (0.005 µM for DHNA) 200-fold lower than HPLC (1 µM for DHNA) (49).

Our work also demonstrates a new concept for improving analyte selectivity in whole cells. Whole-cell biosensors offer advantages such as long-term operation and the ability to function in complex environments, but many of them depend on analyte-specific transcriptional factors. This can be problematic when dealing with structural analogs, as many transcriptional factors must be created to detect each individual analyte (50). In contrast to specific sensing, selective sensing can be achieved through the use of nonspecific sensing elements, such as nanoparticles, quantum dots, peptides, or nonspecific transcriptional factors, to simultaneously detect a class of analytes (51–53). However, these sensors must be carefully arranged in a complex array to generate unique “fingerprints” to identify individual components (51–53). Our quinone bioelectronic sensor represents a new type of whole-cell biosensor that offers both selectivity and specificity. It selectively detects a class of 1,4-naphthoquinones and specifically discriminates between structural analogs, such as menadione and DHNA, based on the temporal dynamics of the electronic output signal (Fig 4A). This difference in output signal is a result of the differing kinetics of Ndh2-dependent reduction of these analogs, which can be described using a multivariate model (Fig 5D). Nonetheless, one question that remains from this work is how to interpret the electronic signals to inform each component when multiple unknown quinones are present in a mixed pool. This can be potentially addressed by building a machine learning model to precisely identify individual components (52, 53).

Moreover, the quinone bioelectronic sensor in this study leverages the native Ndh2 of *L. plantarum* and shows a preference for sensing 1,4-naphthoquinones (Fig. 4B, S6), whereas Ndh2 homologs in other organisms show broader binding to ubiquinones or menaquinones (40, 41). It is possible to use protein evolution to modify Ndh2 of *L. plantarum* to sense other types of quinone derivatives, such as quinone-derived cancer drugs or air pollutants, for rapid drug monitoring and pollution control. Hence, this work opens the possibility of using *L. plantarum* as a probiotic chassis for EET-based bioelectronic applications in the fields of environment, food, and medicine.

## Materials and methods

### Strains and culture conditions

All strains used in this study are listed in Table S4. *Escherichia coli* 10-beta (New England Biolabs) was used for plasmid construction. *Lactiplantibacillus plantarum* NCIMB8826 was used as the parental strain for all the experiments. *L. plantarum* strains were grown in commercial MRS (HiMedia) from glycerol stocks at 37 °C without shaking. To grow a large volume of *L. plantarum* for a subsequent iron reduction assay or a bioelectrochemical assay, cells were subcultured in modified MRS containing 1% (w/v) mannitol and grown at 37 °C without shaking (20, 54). In bioelectrochemical systems, Chemically Defined Medium (CDM) containing 1% (w/v) mannitol (Table S2) was used as the culture medium for *L. plantarum*. For testing in a bio-relevant environment, the Fasted State Simulated Intestinal Fluid (FaSSIF, Biorelevant, UK) was prepared as indicated in Table S3. For both iron reduction and bioelectrochemical assays, cells were tested at 30 °C under anaerobic conditions.

### L. plantarum mutant construction

*L. plantarum* SL046 (Δ*dmkA*Δ*ndh1*) and SL052 (Δ*dmkA*Δ*ndh1*Δ*ndh2*) strains were constructed by using the CRISPR-Cas9 toolbox developed by Huang et al. (55). The crRNAs targeting each gene were designed by using CRISPOR (http://crispor.tefor.net/). The sgRNA fragment and the homologous arms flanking each gene were cloned into the ApaI-XbaI digested pHSP02 backbone by Gibson assembly to create the editing plasmids pSTS04, pSL47, pSL51 that target *dmkA*, *ndh1*, *ndh2*, respectively (primers and oligos are listed in Table S6). For CRISPR editing, *L. plantarum* NCIMB8826 strain harboring helper plasmid pLH01 was induced with 100 ng/ml Sakain P peptide (GenScript) for RecE/T expression and was subsequently prepared as competent cells. The editing plasmid was then delivered into *L. plantarum* by electroporation. The transformed cells were spread on MRS plates containing 10 μg/mL erythromycin and 10 μg/mL chloramphenicol to screen for the deletion mutants. We found that the edited *L. plantarum* mutant does not retain the helper plasmid pLH01 and cannot grow with antibiotic selection in liquid MRS culture. Thus, for sequential editing, we inoculated the *L. plantarum* mutant in antibiotics-free MRS for two passages and screened for the colonies with both the helper plasmid and the editing plasmid being cured. Then, we electroporated the helper plasmid back to the mutant strain for the next round of gene deletion.

### Michaelis-Menten kinetics

We hypothesized that DHNA reduction by Ndh2 follows Michaelis-Menten kinetics when the intracellular NADH is sufficient to support EET. DHNA oxidation by terminal electron acceptor can be either direct or indirect (involving intermediate steps via PplA and/or EetA) (21). When an insoluble terminal electron acceptor is used (such as metal iron or electrode), and the DHNA oxidation is fast enough, this step can be treated as a pseudo-first-order reaction. Under this assumption, the apparent EET rate (V_EET_, the rate that terminal electron acceptors are reduced) is equal to DHNA oxidation rate following first-order kinetics. At the quasi-steady-state, the concentration of reduced DHNA does not change. This means DHNA reduction rate is equal to DHNA oxidation rate:

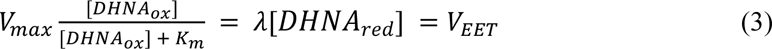

thus,

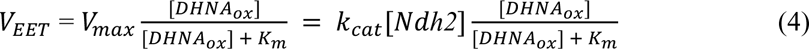

where λ is the first-order rate constant, V_max_ is the maximum DHNA reduction rate, K_m_ is the Michaelis constant, k_cat_ is the turnover number of Ndh2. This implies that the apparent EET rate (V_EET_) is a Michaelis-Menten function of oxidized DHNA concentration. Because it is unpractical to measure the concentration of oxidized DHNA at steady-state, we assume [DHNA_ox_] ≫ [DHNA_red_] when DHNA oxidation is fast enough and treat the total DHNA concentration [DHNA_total_] as an approximation of the oxidized DHNA concentration:

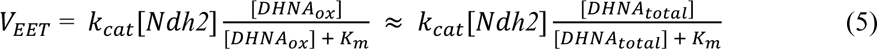

Therefore, we used equation 5 to fit the experimental data into Michaelis-Menten equation. All data fitting and analysis were performed in either OriginLab or MATLAB.

### Iron reduction assay

After overnight growth, cells were harvested at 4000 rpm, 4 °C, for 15 min. Cells were washed twice with 1x PBS and resuspended in 1x PBS to OD_600_=4. CFU was then measured by dilution plating. In an anaerobic chamber (Whitley A45 anaerobic workstation), the concentrated cells were inoculated in a 96-deep well plate (cells were series diluted if necessary) and mixed in a 1:1 ratio with a 2x iron master mixture prepared in PBS. The final composition of the reaction mixture was: 2 mM iron(III) oxide nanoparticles (< 50 nm, Sigma-Aldrich), 2 mM ferrozine, 1% (m/v) mannitol, indicated concentration of DHNA, and indicated cell OD_600_. After incubating at 30 °C for indicated hours, a 150 μl aliquot of the reaction mixture was harvested, and 100 μl supernatant was collected after centrifuging at 4000 rpm for 10 min. The absorbance of the supernatant was measured at 562 nm by a plate reader (Tecan Spark). The concentration of Fe^2+^ in each sample was calculated by a standard curve prepared with FeSO_4_ (0-0.4 mM, 2-fold increase) and was normalized to CFU. The scanned image showing the color range was taken by a desktop scanner in a 96-well white bottom plate.

### Bioelectrochemical system construction

We used water-jacketed dual-chamber bioelectrochemical systems (BESs) (Adams & Chittenden Scientific Glass) and a VMP-300 potentiostat (BioLogic) for all the bioelectrochemical measurements. The anodic chamber of the BES contained an Ag/AgCl reference electrode filled with 3M KCl (CH111, CH Instruments) and a 6.35-mm-thick graphite felt working electrode with a 16-mm radius size (Alfa Aesar) threaded to a 0.5-mm radius titanium wire (Alfa Aesar). The cathodic chamber contained a titanium wire as the counter electrode. The anodic chamber was separated from the cathodic chamber by using a cation membrane (CMI-7000, Membranes International).

The BESs were filled with ddH_2_O and sterilized by autoclaving. The media in the anodic chamber was then replaced by 110 ml 1x CDM or PBS containing 1% mannitol. The media in the cathodic chamber was replaced by 1x M9 salts (BD Difico). A magnetic stir bar was placed in the anodic chamber for continuous stirring at 220 rpm (IKA RO10 Magnetic Stirrers). N_2_ gas was purged continuously into the anodic chamber to maintain anaerobic conditions. The BESs were kept at 30 ℃ by connecting the water-jackets to an ECO ES4 heating circulator (Lauda-Brinkmann). To carry out bioelectrochemical measurements, the working electrode was poised at +0.2 V versus Ag/AgCl and chronoamperometry was carried out to record the current every 36 s. Once the current stabilized, the cells were inoculated for the subsequent bioelectrochemical analysis.

### Bioelectrochemical assay with quinone addition

Overnight cultures of *L. plantarum* were harvested at 4000 rpm, 4 ℃, for 15 min. Cells were washed twice and resuspended in 1x PBS to OD_600_ = 9.16 per ml. A 3 ml aliquot of the cell resuspension was injected into the BESs so that the final OD_600_ = 0.25 per ml. To measure CFU, a 200 μl aliquot of sample was taken from each BESs after cells were homogeneously stirred. The CFU was then determined by dilution plating. The quinone of interest was injected after the current had stabilized.

DHNA, hydroquinone, 1,4-naphthoquinone, juglone, DCDMNQ, menadione, and lawsone (Sigma-Aldrich) were prepared freshly in 100% dimethyl sulfoxide (DMSO). A 200 μl aliquot of a 550x stock quinone solution with varied concentrations was then injected into the BESs containing 110 ml medium for the desired final concentration (1x). The current was recorded every 1s to monitor the transient current change upon quinone addition.

### Redox potential measurement or prediction

Cyclic voltammetry (CV) measurement of quinone 1e^-^ or 2e^-^/2H^+^ redox potential was conducted in a single chamber electrochemical cell with a glassy carbon working electrode and a Pt mesh counter electrode. All solutions were sparged with N_2_ prior to the measurement and N_2_ was continuously blown over the solution during the CV scan to maintain an anaerobic condition. The 2e^-^/2H^+^ redox potential was measured in 100 mM MOPS buffer containing 100 μM quinone of interest. CV was performed v.s. Saturated Calomel Electrode (SCE) reference electrode at a scan rate of 50 mV/s. The 1e^-^ redox potential was measured in acetonitrile containing 0.1 M tetrabutylammonium hexafluorophosphate (TBAPF6) as the electrolyte (56). Each quinone of interest was tested at 10 mM. Ag/Ag^+^ was used as the reference electrode. Ferrocene of 5 mM was added as the internal standard. CV was performed at a scan rate of 50 mV/s.

The 1e^-^ redox potential for DHNA was not experimentally accessible due to the lack of purified oxidized DHNA. We adapted and modified the method reported by Prince et al. to correlate the Hammett σ_para_ constants with 1e^-^ redox potentials (45). We then used the trend line to predict the 1e^-^ redox potential for DHNA (see Fig. S11).

### Quinone-Ndh2 binding simulation

The three-dimensional structure of Ndh2 from *L. plantarum* was predicted by AlphaFold (39). The truncated 1-409 aa region was used for molecular docking (Fig. S9). Quinone-Ndh2 docking was performed by using AutoDock Vina (37) with the searching exhaustiveness of 9 and the energy range of 24 (kcal/mol). The grid box size was set to 24Å × 34Å × 10Å and centered at position (8.885, −7.046, 6.636) in the coordinate space of the receptor. This space is demonstrated to be the quinone binding pocket according to previous studies (40, 41). The model with the highest binding energy and a proper posing of quinone in the binding pocket was chosen for further analysis.

### cLogP determination

The cLogP value for quinone or semiquinone was calculated by using ChemDraw 21.0.0.

## Acknowledgments

We thank Dr. Jayashree Soman (Rice University) for helping predict the Ndh2 structure and reviewing the manuscripts; Prof. Emily Mevers (Virginia Tech) for discussing the quinone-EET relationship; Prof. Maria Marco, Dr. Eric Stevens (University of California, Davis), and Dr. Sara Tejedor-Sanz (Lawrence Berkeley National Laboratory) for helpful conversations; Prof. Shannon Stahl and Dr. James Gerken (University of Wisconsin–Madison) for the protocol of measuring quinone 1e^-^ redox potential. This work was supported by the Cancer Prevention and Research Institute of Texas awarded # RR190063 and by the Army Research Office awarded W911NF-22- 1-0239.

## Supplementary figures

**Fig S1.**
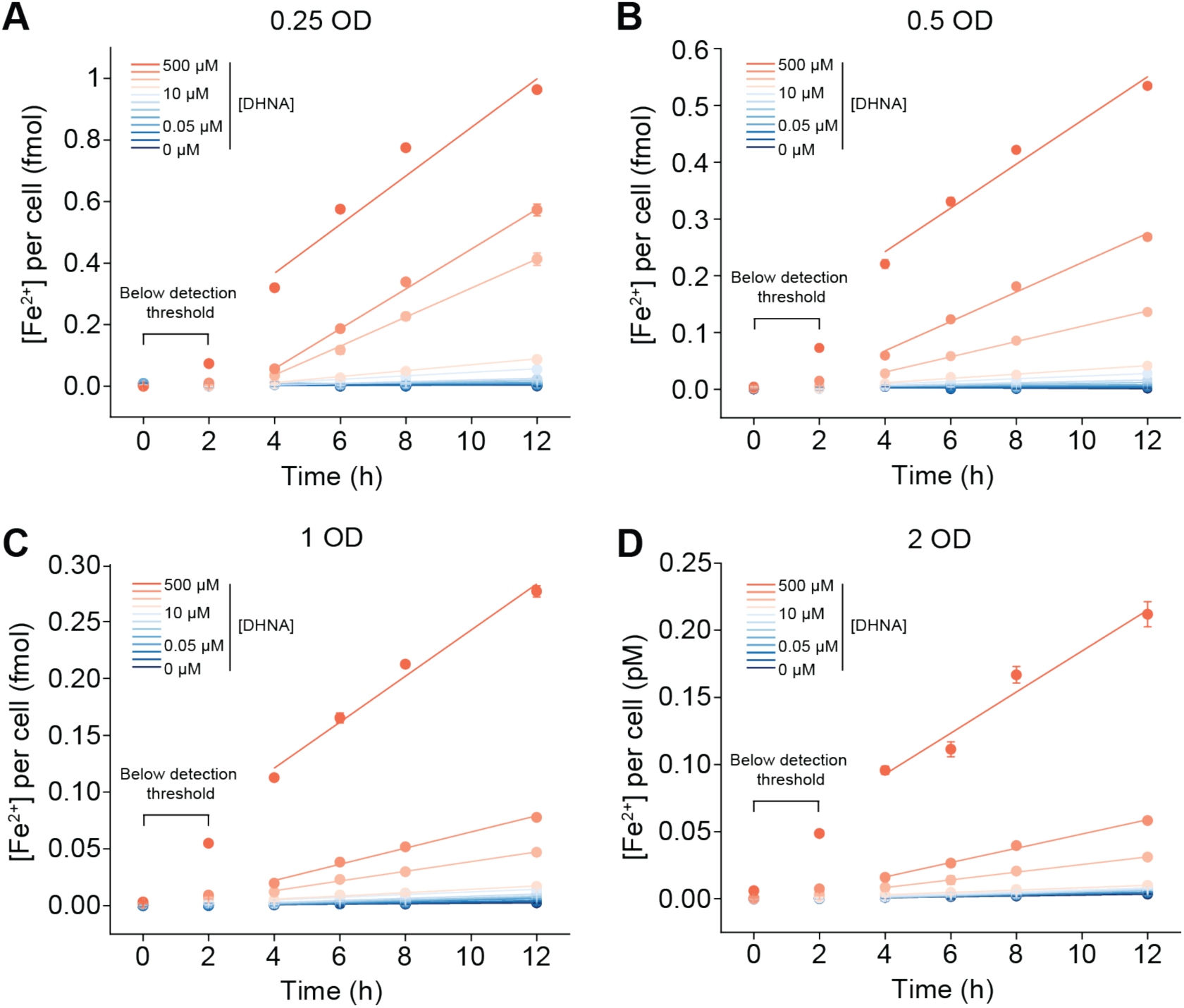
Time-course iron reduction and calculation of the initial iron reduction rate. (A-D) Concentration of Fe^2+^ produced by 0.25, 0.5, 1, or 2 OD of cells was measured at 0, 2, 4, 6, 8, 12 h after cells were mixed with iron(III) oxide and different concentrations of DHNA. To calculate the initial iron reduction rate, data of 0 h and 2 h were omitted for all DHNA concentrations and for all cell densities to improve the accuracy of linear regression. This is because the Fe^2+^ concentrations were below the detection threshold at these two time points. The rest of the data were fitted by linear regression and the slope value was treated as the iron reduction rate. Data represent mean ± 1 s.d. for n=3 biological replicates.

**Fig S2.**
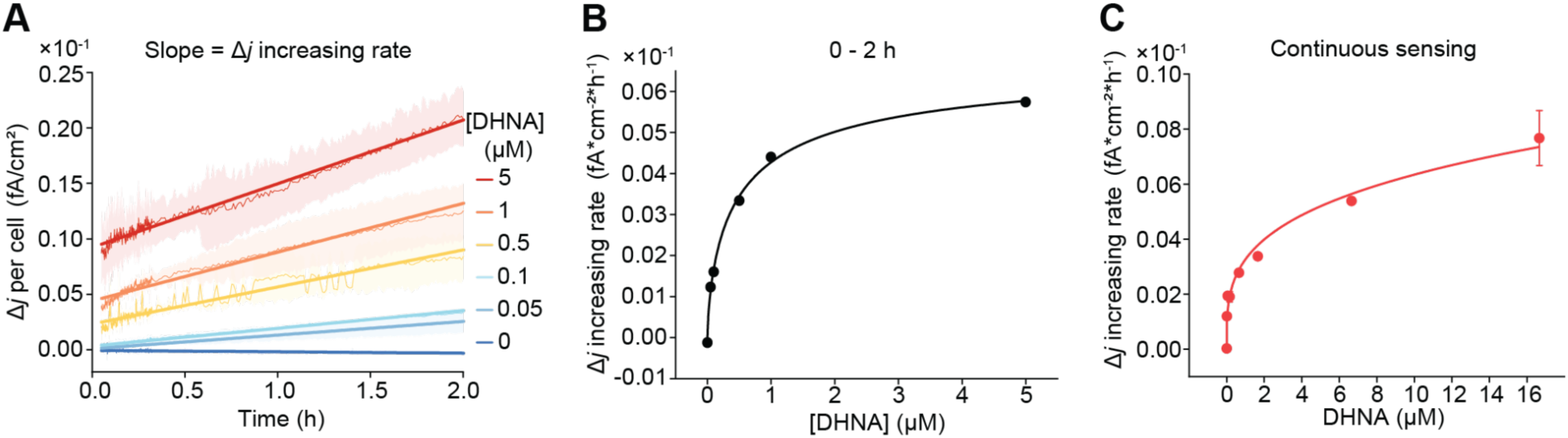
Correlation between DHNA concentration and the increasing rate of current density. (A) The current density produced by *L. plantarum* Δ*dmkA*Δ*ndh1* strain within 0 - 2 h in response to different concentrations of DHNA (based on the data shown in Figure 3) was fitted by linear regression. The solid lines represent the result of linear regression. The slope value indicates the increasing rate of current density. (B) The increasing rate of current density as a function of DHNA concentrations. The line is a guide to the eye. (C) In the DHNA continuous sensing experiment shown in Figure 3D, the linear regression of current density was performed on each adding interval. The increasing rate of current density was plotted against the accumulated DHNA concentration. The line is a guide to the eye. Data represent mean ± 1 s.d. for n=2 biological replicates.

**Fig S3.**
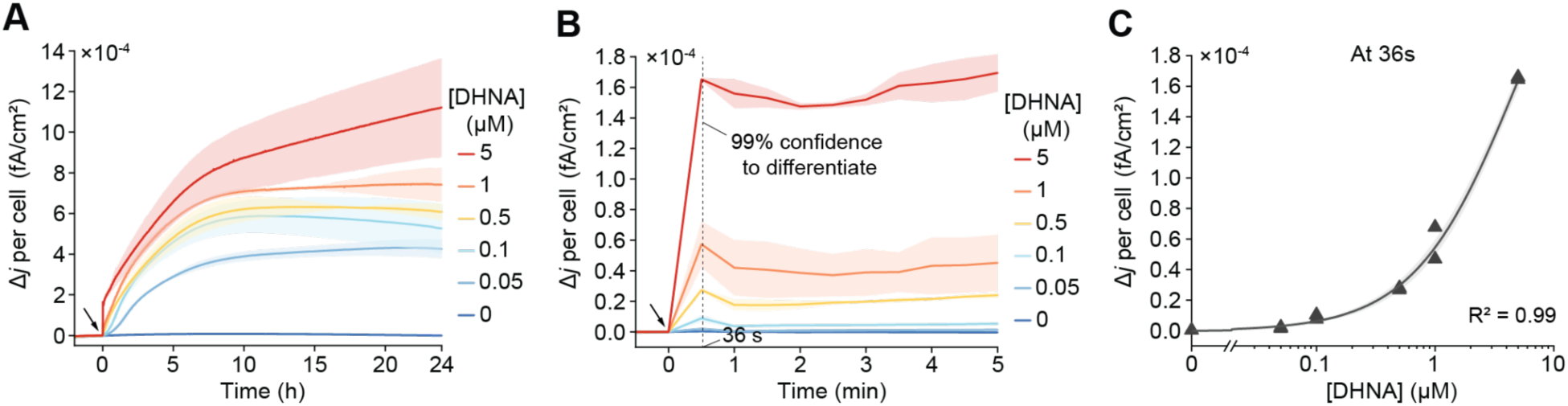
*L. plantarum* responds to DHNA in PBS. (A) Current density produced by *L. plantarum* Δ*dmkA*Δ*ndh1* strain upon exposure to different concentrations of DHNA with PBS as the assay buffer. The black arrow indicates DHNA injection. Data represent mean ± 1 s.d. for n=2 biological replicates. (B) An enlarged region of (A) shows a rapid current response within 5 minutes upon exposure to environmental DHNA in PBS. The dash black line indicates the time with 99% confidence (p-value ≤ 0.001) to discriminate different DHNA concentrations. (C) Current density as a function of DHNA concentration at the time when 99% confidence is reached. Data were fitted in the Michaelis-Menten equation.

**Fig S4.**
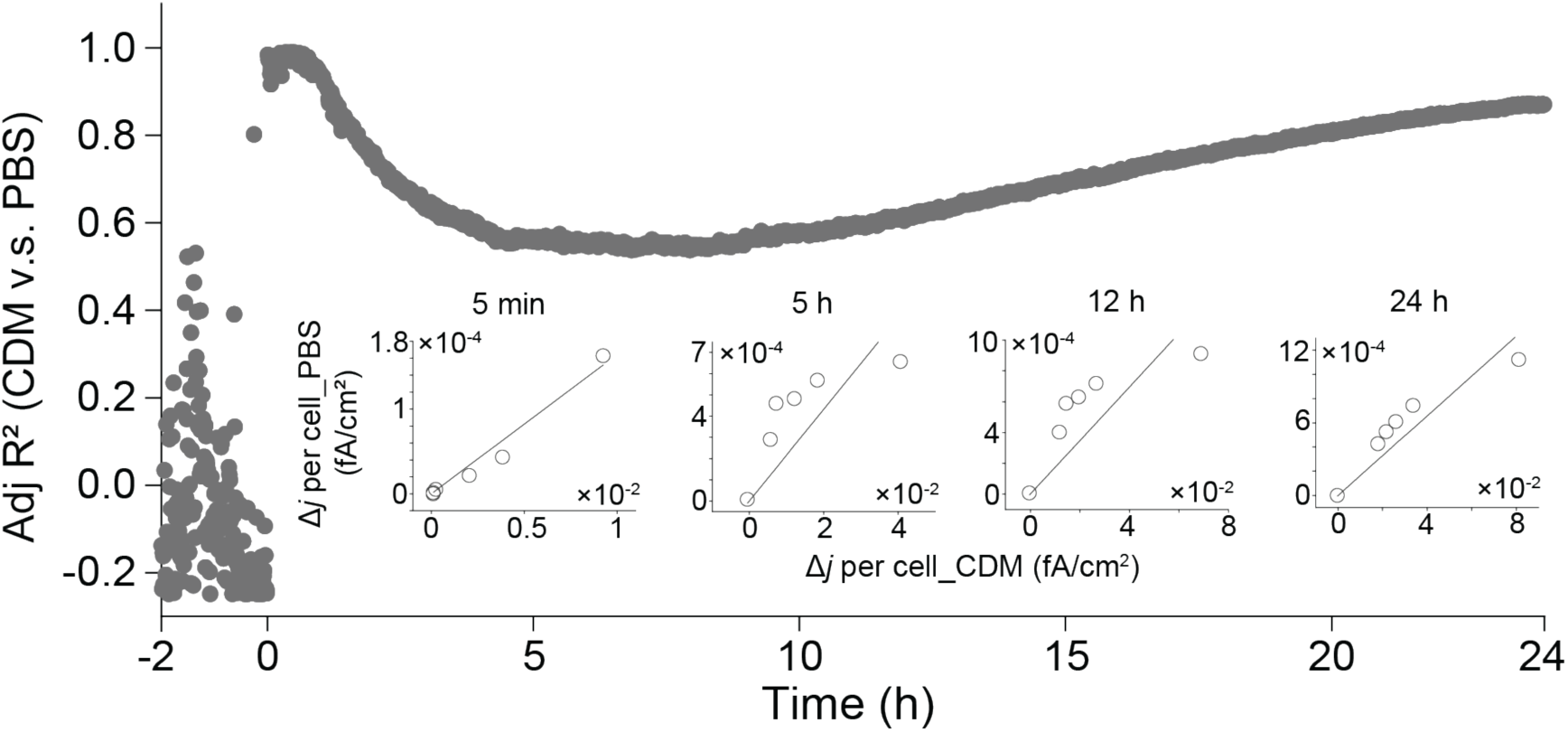
Comparison of current density produced by *L. plantarum* in response to DHNA in different media. Current densities at each time point were compared between CDM and PBS and the adjusted R^2^ was reported. Insets show the comparison at 5 min, 5 h, 12 h, and 24 h. Each dot in the insets represent a different DHNA concentration. The mean data of n=2 biological replicates for both CDM and PBS were used for comparison.

**Fig S5.**
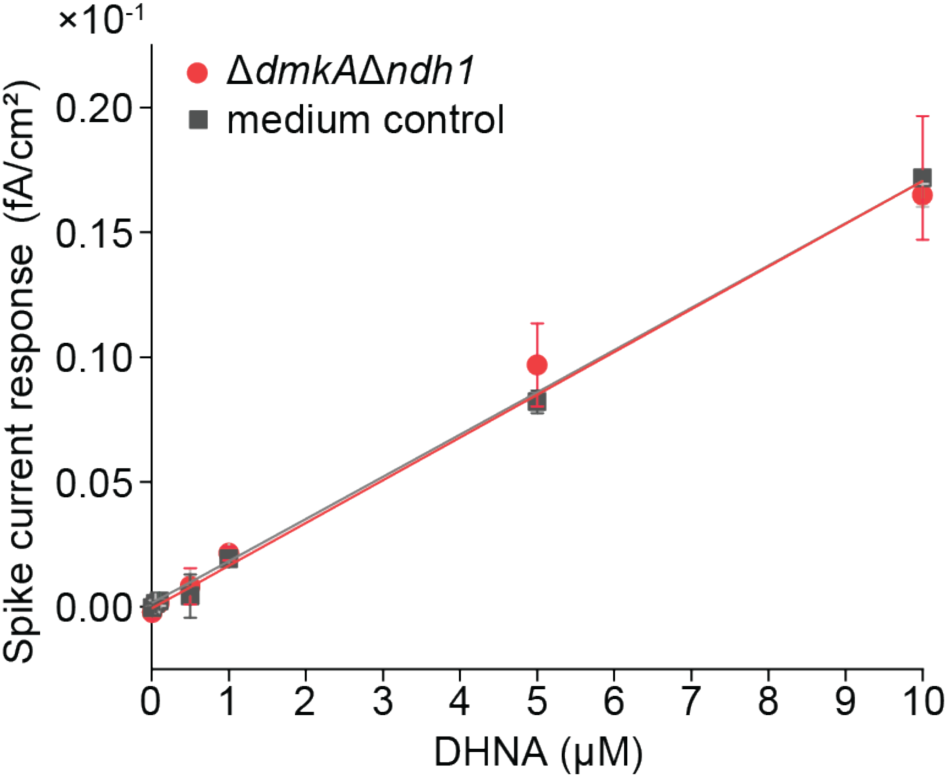
The current spike upon DHNA addition results from abiotic DHNA oxidation. The amplitude of the current spike upon DHNA injection was calculated for the Δ*dmkA*Δ*ndh1* mutant and the medium control. The spike current density was plotted against DHNA concentrations. No difference was observed between these two groups. Data represent mean ± 1 s.d. for n=2 biological replicates.

**Fig S6.**
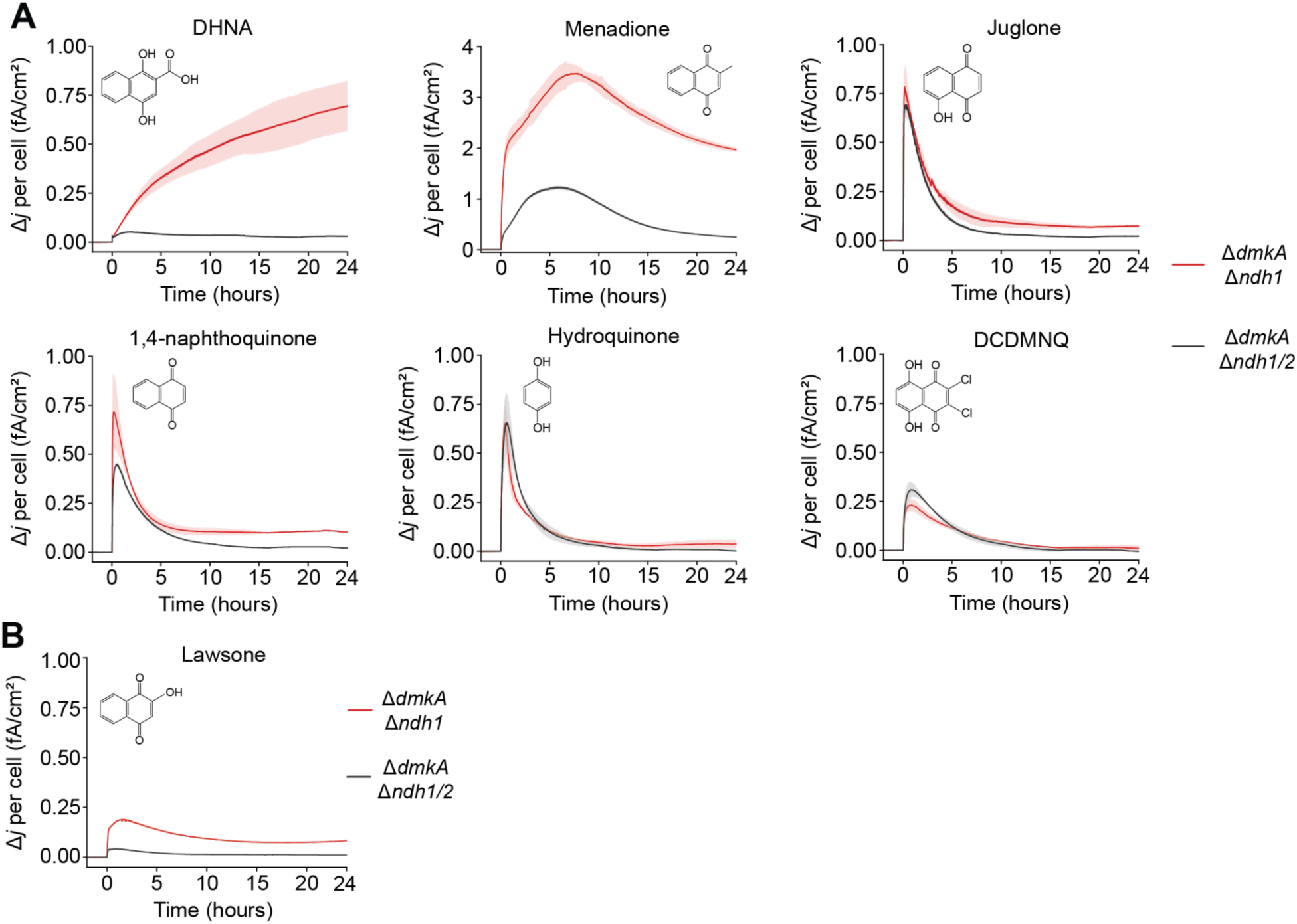
Current density generated by *L. plantarum* in response to different quinone derivations. (A) The indicated quinone derivatives (1 μM) were injected at 0 h to either Δ*dmkA*Δ*ndh1* or Δ*dmkA*Δ*ndh1/2* strain. The two mutant strains produced differentiated current density when reacting with DHNA or menadione but not with other quinone derivatives. Data represent mean ± 1 s.d. for n=2 biological replicates. (B) Lawsone can be utilized by Ndh2- dependent EET to produce current, in agreement with the model’s prediction shown in Fig. 5E. Data represent mean ± 1 s.d. for n=2 biological replicates.

**Fig 7.**
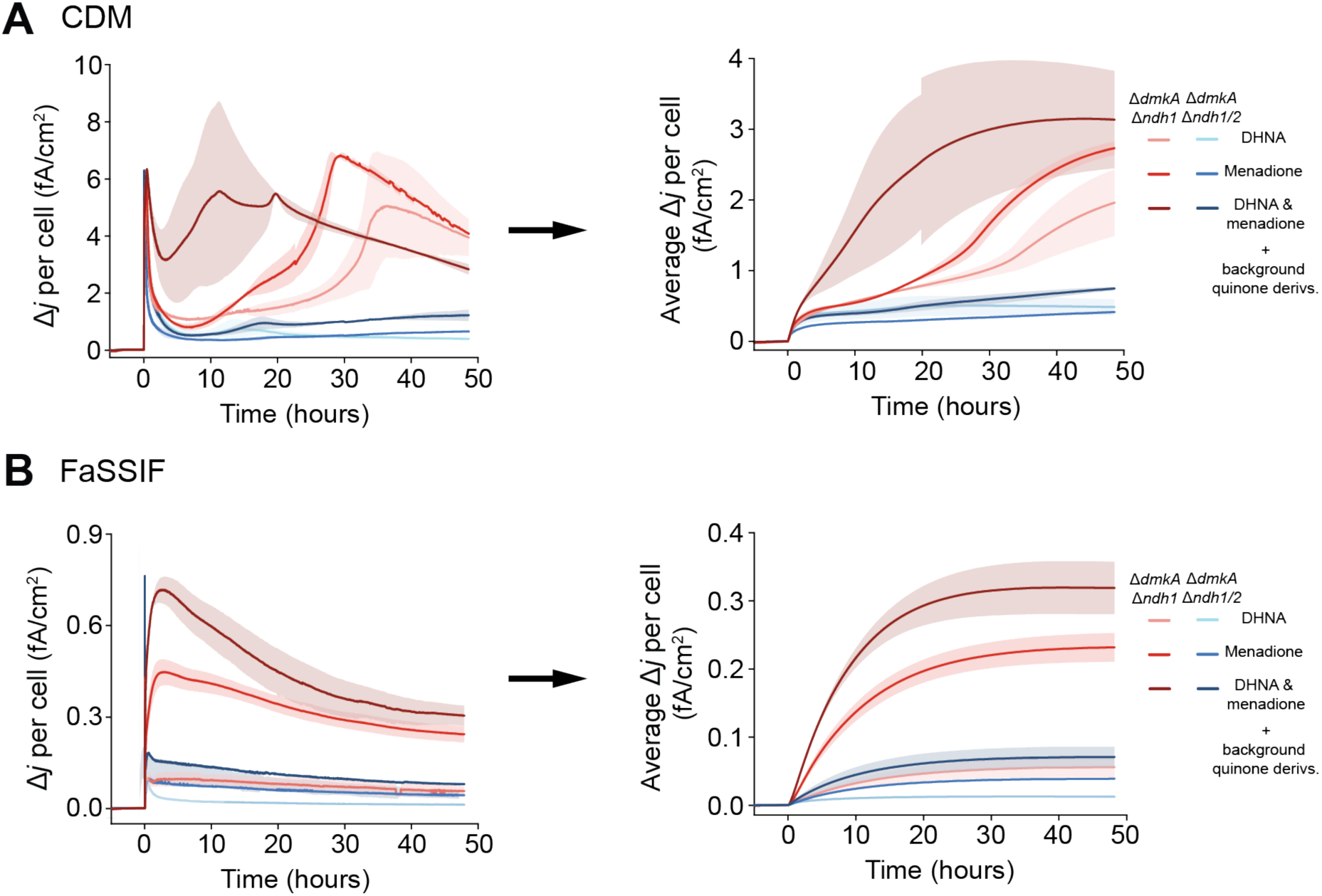
Average current density is less dependent of media than current density, allowing data comparison between different media. (A) Data transition from current density (dQ/dt per cm^2^) to average current density (ΔQ/Δt per cm^2^) in CDM. (B) Data transition from current density to average current density in FaSSIF. Data represent mean ± 1 s.d. for n=2 biological replicates.

**Fig S8.**
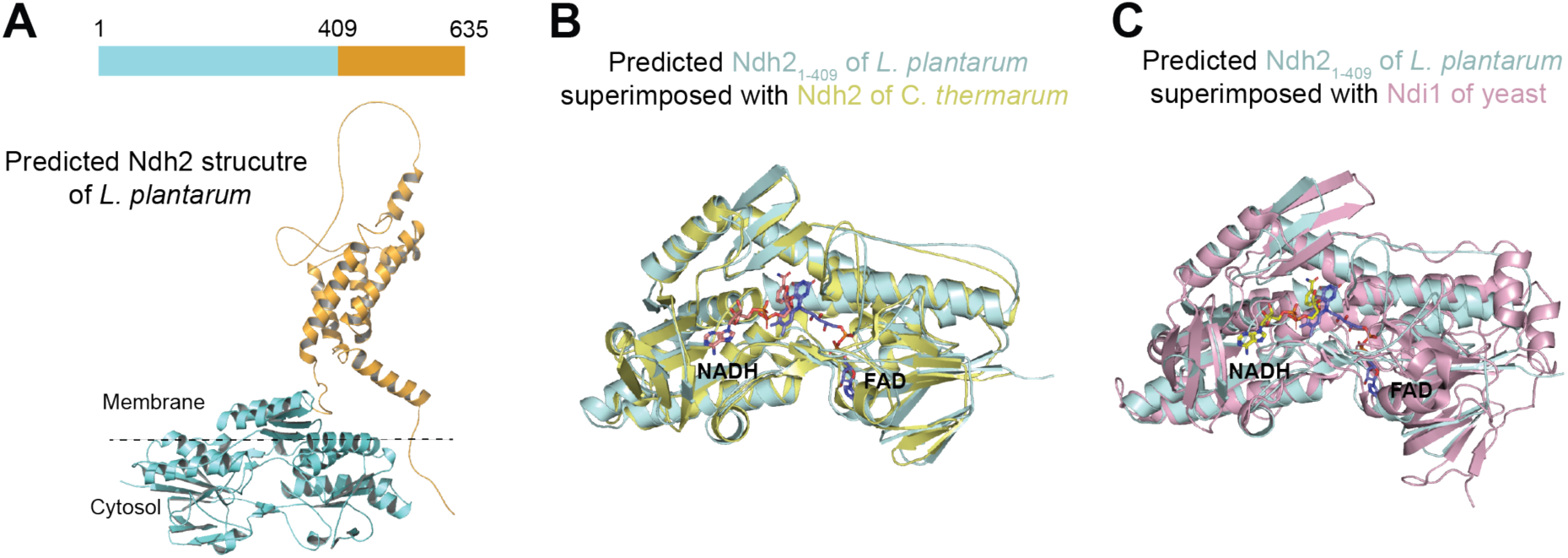
Ndh2 structure prediction and comparison. (A) AlphaFold predicted Ndh2 structure of *L. plantarum*. The cyan region (1-409 aa) shows homology with known crystal structures of Ndh2. The orange region (409-635 aa) shows no homology to known protein. Truncated Ndh2_1-409_ was used for structural comparison and for AutoDock Vina simulation. (B) Truncated Ndh2_1-409_ of *L. plantarum* superimposed upon Ndh2 of *Caldalkalibacillus thermarum* (PDB: 5KMS). (C) Truncated Ndh2_1-409_ of *L. plantarum* superimposed upon Ndi1 of yeast (PDB: 4G73).

**Fig S9.**
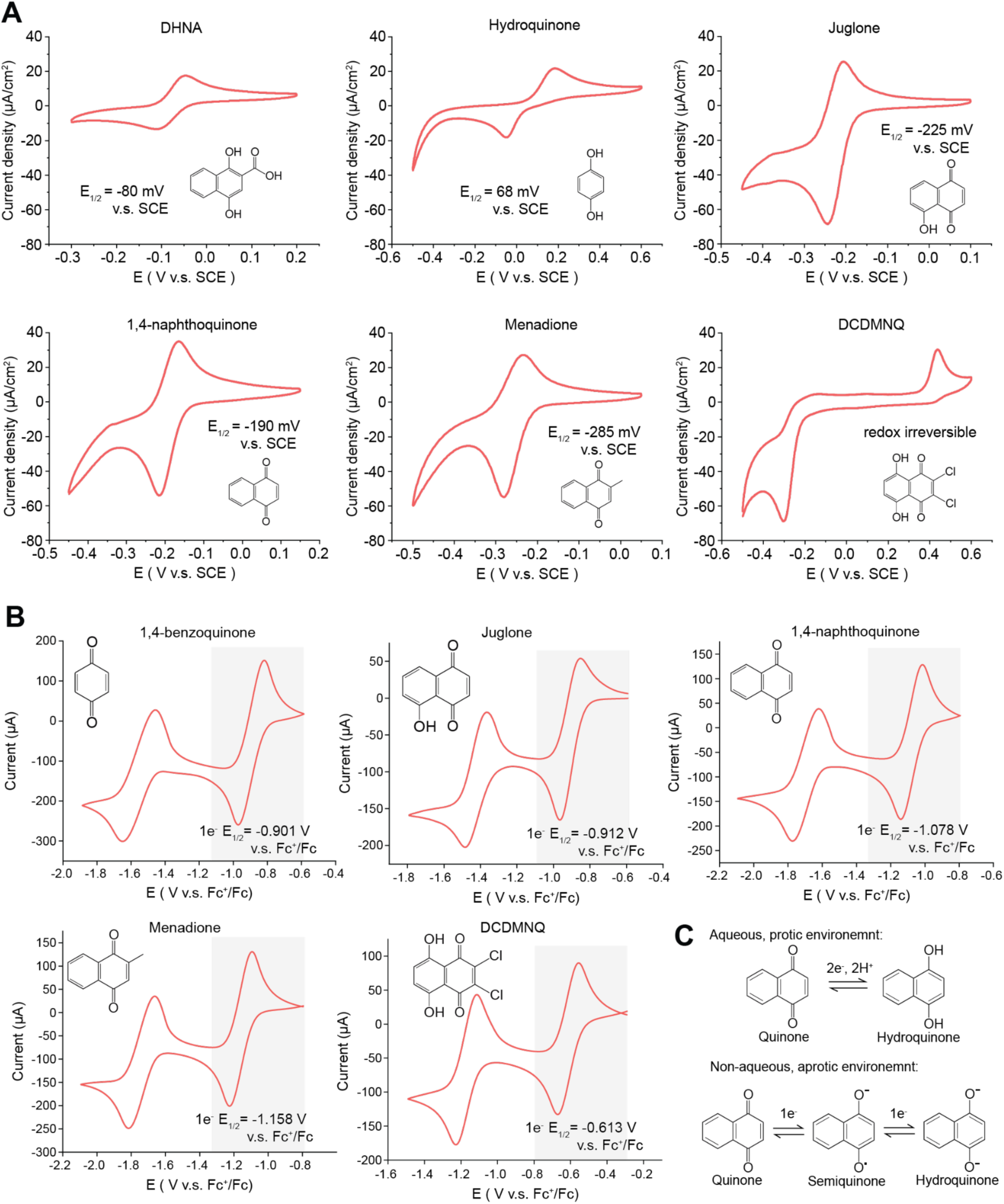
Cyclic voltammetry measurements of quinone 1 e^-^ or 2 e^-^/ 2 H^+^ redox potential. (A) The 2 e^-^/ 2 H^+^ redox potential measurements of the represented quinone derivatives in an aqueous solution (MOPS buffer). DCDMNQ is redox irreversible in the aqueous solution despite the two well-separated redox peaks. (B) Quinone 1 e^-^ redox potential measurements in a non-aqueous solution (acetonitrile). Gray shadows indicate the redox peaks corresponding to 1 e^-^ transfer. (C) Quinone undergoes 2 e^-^/2 H^+^ electron transfer in aqueous environment and successive 1 e^-^ electron transfer in non-aqueous environment.

**Fig S10.**
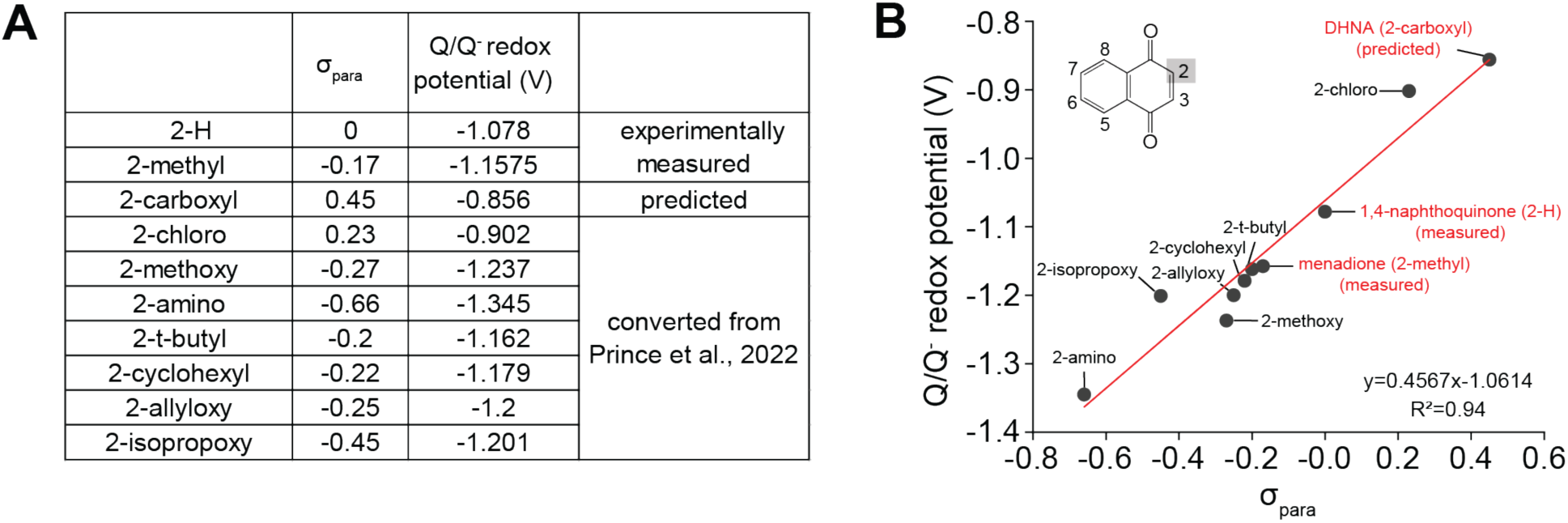
Prediction of the 1 e^-^ redox potential of DHNA. (A) Summary of Hammett σ_para_ constants for each 2-substitutions of 1,4-naphthoquinone and the 1e^-^ redox potentials for each derivatives. The 1e^-^ redox potential for 1,4-naphthoquinone (2-H) and menadione (2-methyl) were measured in this paper (Fig. S9). The 1e^-^ redox potentials for other derivatives were reported by Prince et al. and were converted to compare with our measurements by a factor of −0.5 V. The Hammett σ_para_ constants were then plotted against 1e^-^ redox potentials and the 1e^-^ redox potential of DHNA (2-carboxyl) was predicted by using the fitted equation. (B) Hammett σ_para_ constants plotted against 1e^-^ redox potential of 2-substituted 1,4-naphthoquinones showing a linear relationship between these two parameters.

**Fig S11.**
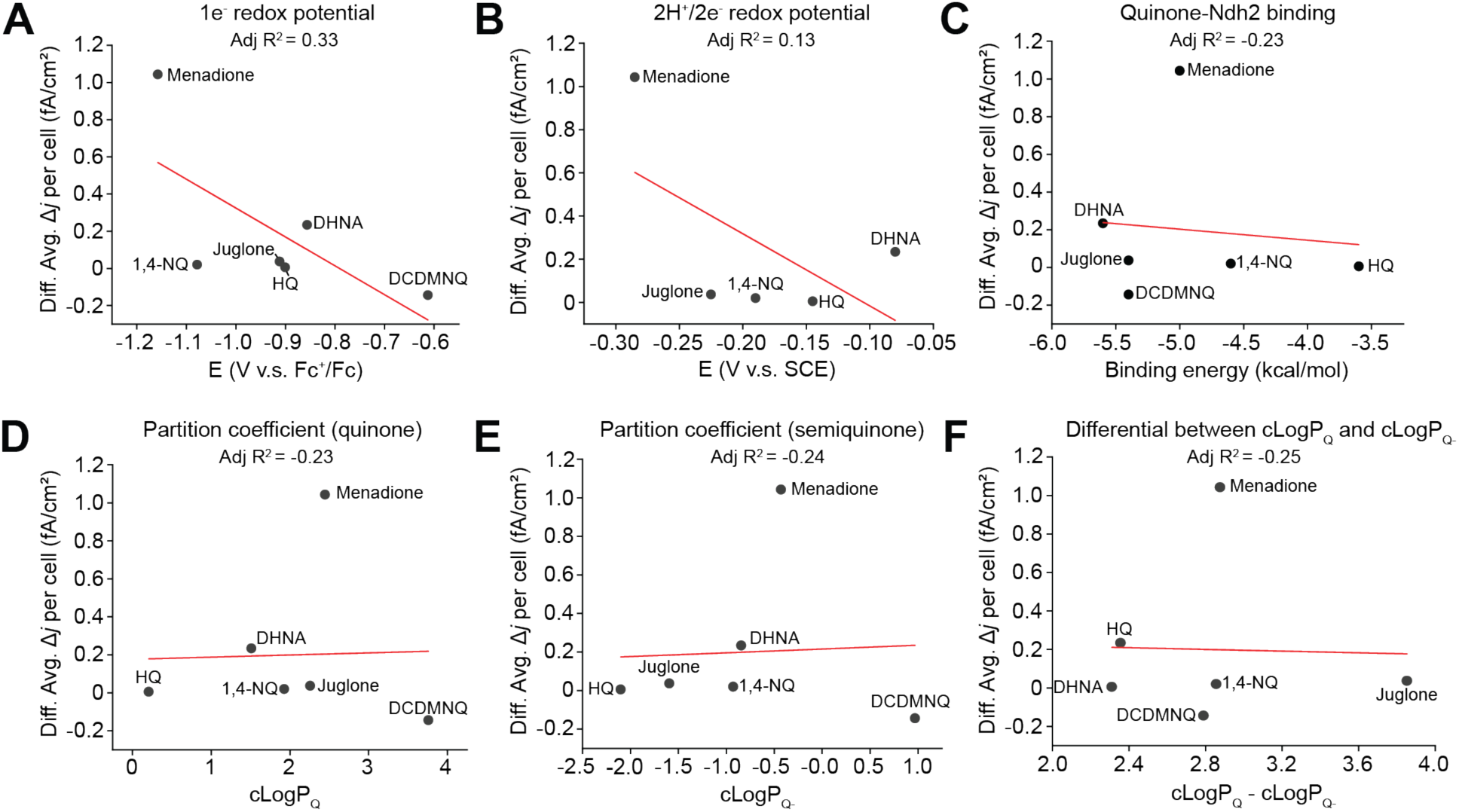
Relationship between EET and quinone physiochemical properties. Differential average current density plotted against (A) quinone one electron redox potential, (B) quinone two electrons redox potential, (C) Autodock Vina predicted quinone-Ndh2 binding energy, (D) calculated partition coefficient of quinone (cLogP_Q_), (E) calculated partition coefficient of semiquinone (cLogP_Q-_), (F) differentail of the cLogP between quinone and semiquinone. NQ, naphthoquinone; HQ, hydroquinone.

## Supplementary Tables

**Table S1:**
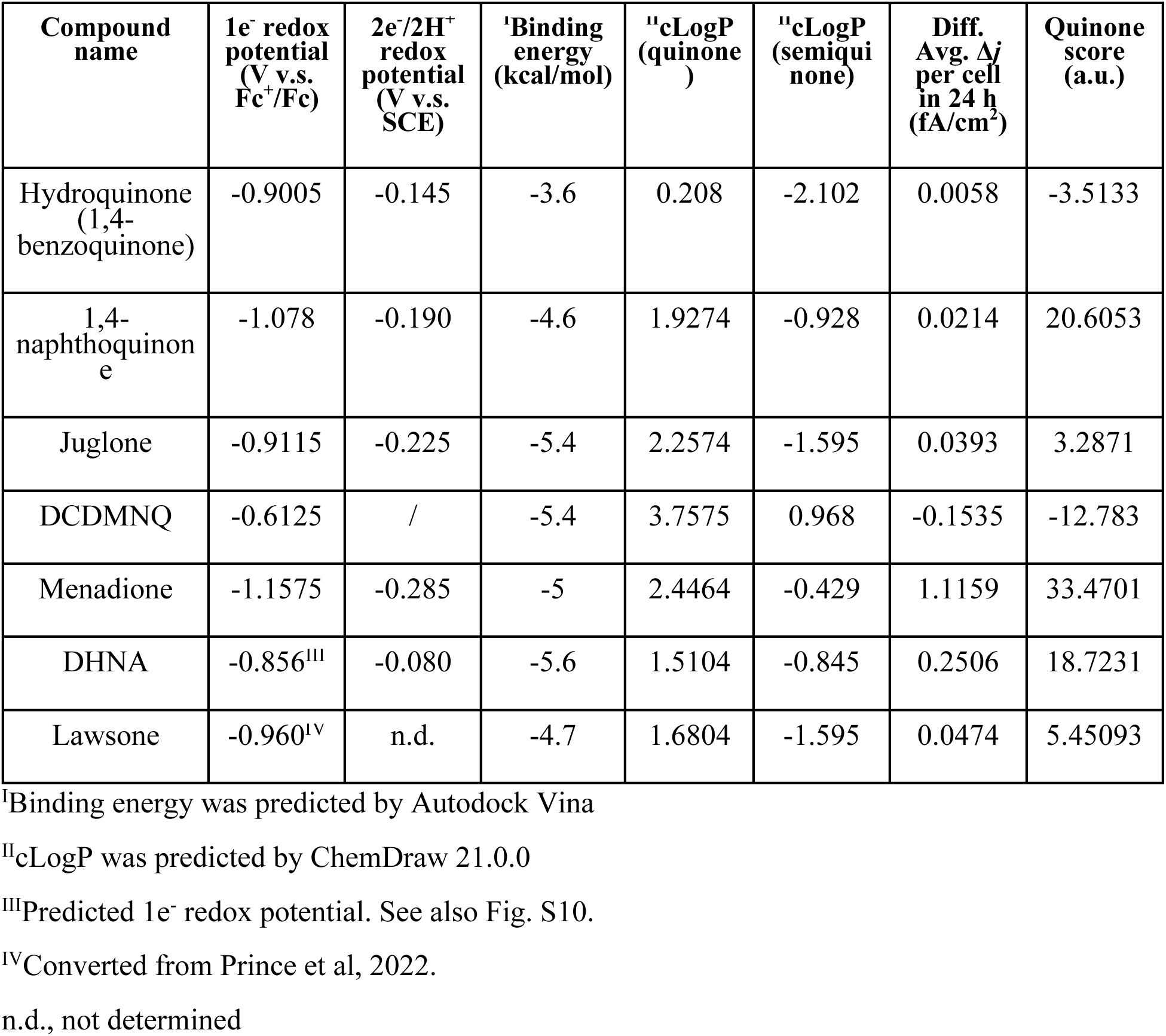
Quinone parameters summary.

**Table S2:**
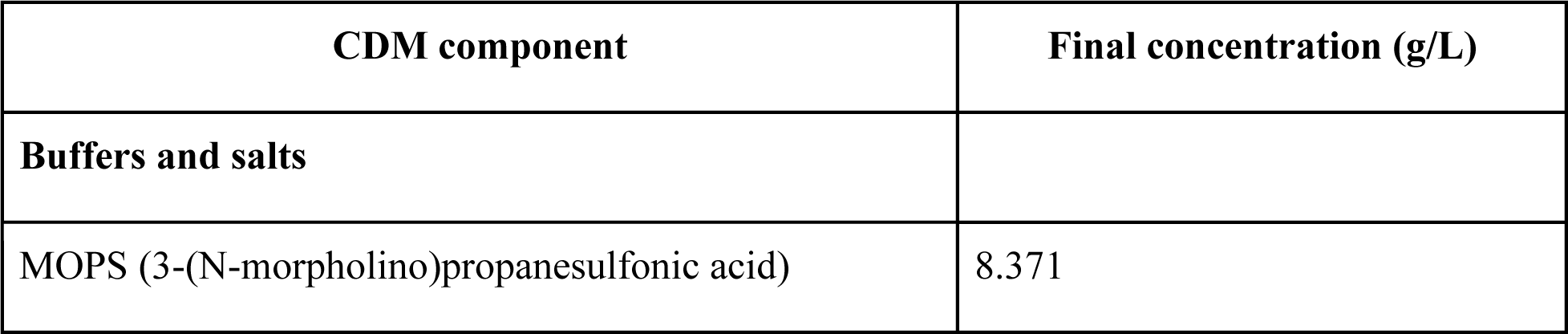
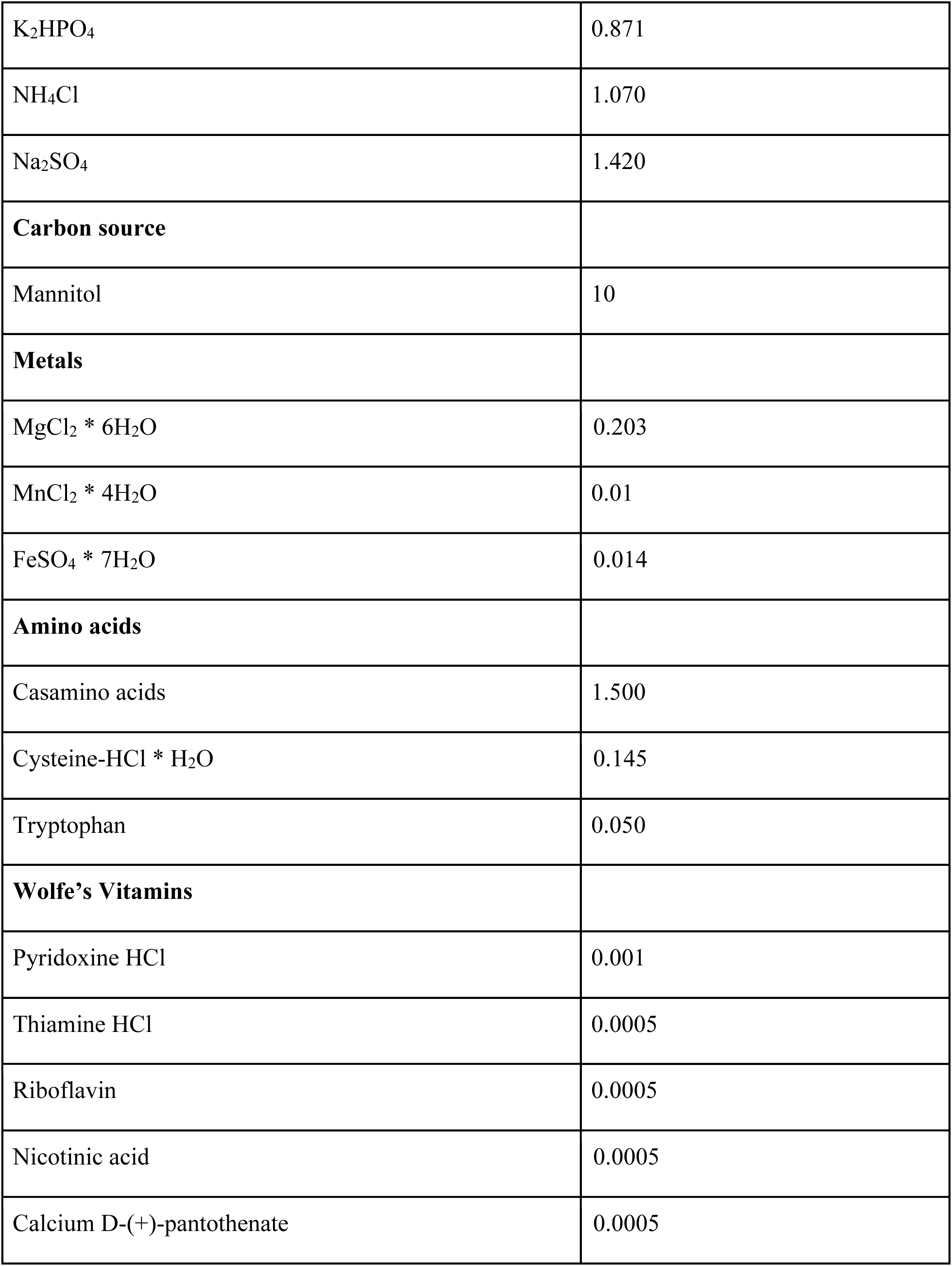
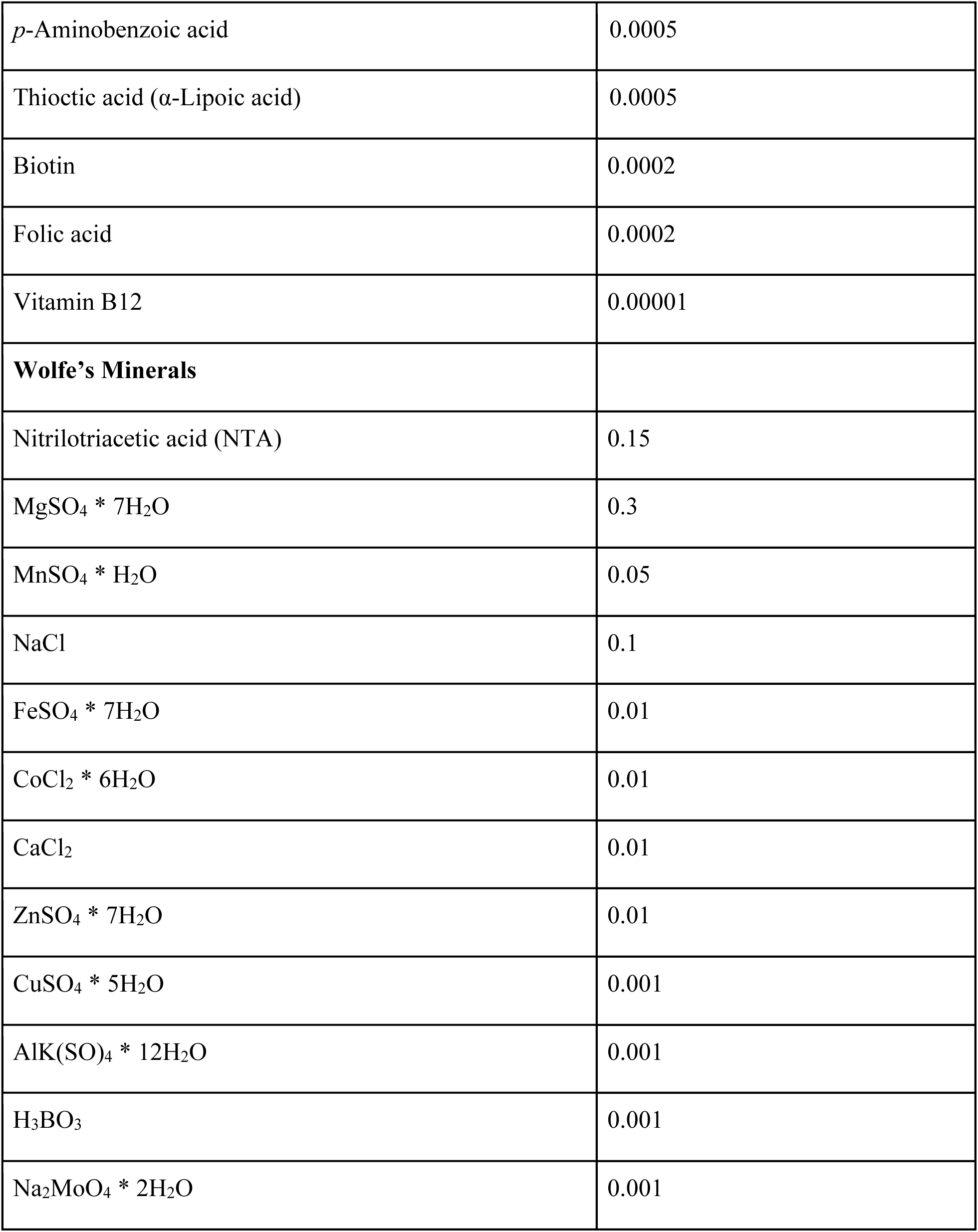
Chemical defined medium recipe.

| CDM component | Final concentration (g/L) |
| --- | --- |
| **Buffers and salts** |  |
| MOPS (3-(N-morpholino)propanesulfonic acid) | 8.371 |
| K<sub>2</sub>HPO<sub>4</sub> | 0.871 |
| NH<sub>4</sub>Cl | 1.070 |
| Na<sub>2</sub>SO<sub>4</sub> | 1.420 |
| **Carbon source** |  |
| Mannitol | 10 |
| **Metals** |  |
| MgCl<sub>2</sub> * 6H<sub>2</sub>O | 0.203 |
| MnCl<sub>2</sub> * 4H<sub>2</sub>O | 0.01 |
| FeSO<sub>4</sub> * 7H<sub>2</sub>O | 0.014 |
| **Amino acids** |  |
| Casamino acids | 1.500 |
| Cysteine-HCl * H<sub>2</sub>O | 0.145 |
| Tryptophan | 0.050 |
| **Wolfe's Vitamins** |  |
| Pyridoxine HCl | 0.001 |
| Thiamine HCl | 0.0005 |
| Riboflavin | 0.0005 |
| Nicotinic acid | 0.0005 |
| Calcium D-(+)-pantothenate | 0.0005 |
| *p*-Aminobenzoic acid | 0.0005 |
| Thioctic acid (α-Lipoic acid) | 0.0005 |
| Biotin | 0.0002 |
| Folic acid | 0.0002 |
| Vitamin B12 | 0.00001 |
| **Wolfe's Minerals** |  |
| Nitrilotriacetic acid (NTA) | 0.15 |
| MgSO<sub>4</sub> * 7H<sub>2</sub>O | 0.3 |
| MnSO<sub>4</sub> * H<sub>2</sub>O | 0.05 |
| NaCl | 0.1 |
| FeSO<sub>4</sub> * 7H<sub>2</sub>O | 0.01 |
| CoCl<sub>2</sub> * 6H<sub>2</sub>O | 0.01 |
| CaCl<sub>2</sub> | 0.01 |
| ZnSO<sub>4</sub> * 7H<sub>2</sub>O | 0.01 |
| CuSO<sub>4</sub> * 5H<sub>2</sub>O | 0.001 |
| AlK(SO)<sub>4</sub> * 12H<sub>2</sub>O | 0.001 |
| H<sub>3</sub>BO<sub>3</sub> | 0.001 |
| Na<sub>2</sub>MoO<sub>4</sub> * 2H<sub>2</sub>O | 0.001 |

**Table S3:** Fasted state simulated intestinal fluid recipe.

| Components | Final concentration (g/L) |
| --- | --- |
| NaOH | 0.42 |
| NaH<sub>2</sub>PO<sub>4</sub>*H<sub>2</sub>O | 3.95 |
| NaCl | 6.19 |
| FaSSIF powder | 2.24 |

**Table S4:** Strains used in this study.

| Strains | Description | Source |
| --- | --- | --- |
| *E.coli* 10-beta | For molecular cloning | New England Biolabs |
| *L. plantarum* NCIMB8826 | NCIMB8826 wide type | Tejedor-Sanz et al., 2022 |
| *L. plantarum* SL046 | NCIMB8826 $\Delta dmkA\Delta ndh1$ | This study |
| *L. plantarum* SL052 | NCIMB8826 $\Delta dmkA\Delta ndh1\Delta ndh2$ | This study |

**Table S5:** Plasmids used in this study.

| Plasmids | Description | Reference |
| --- | --- | --- |
| pLH01 | Helper plasmid for CRISPR-Cas9 toolbox containing RecE/T recombinase | Huang et al., 2019 |
| pSL08 | Editing plasmid for *dmkA* knockout | This study |
| pSL47 | Editing plasmid for *ndh1* knockout | This study |
| pSL51 | Editing plasmid for *ndh2* knockout | This study |

**Table S6:**
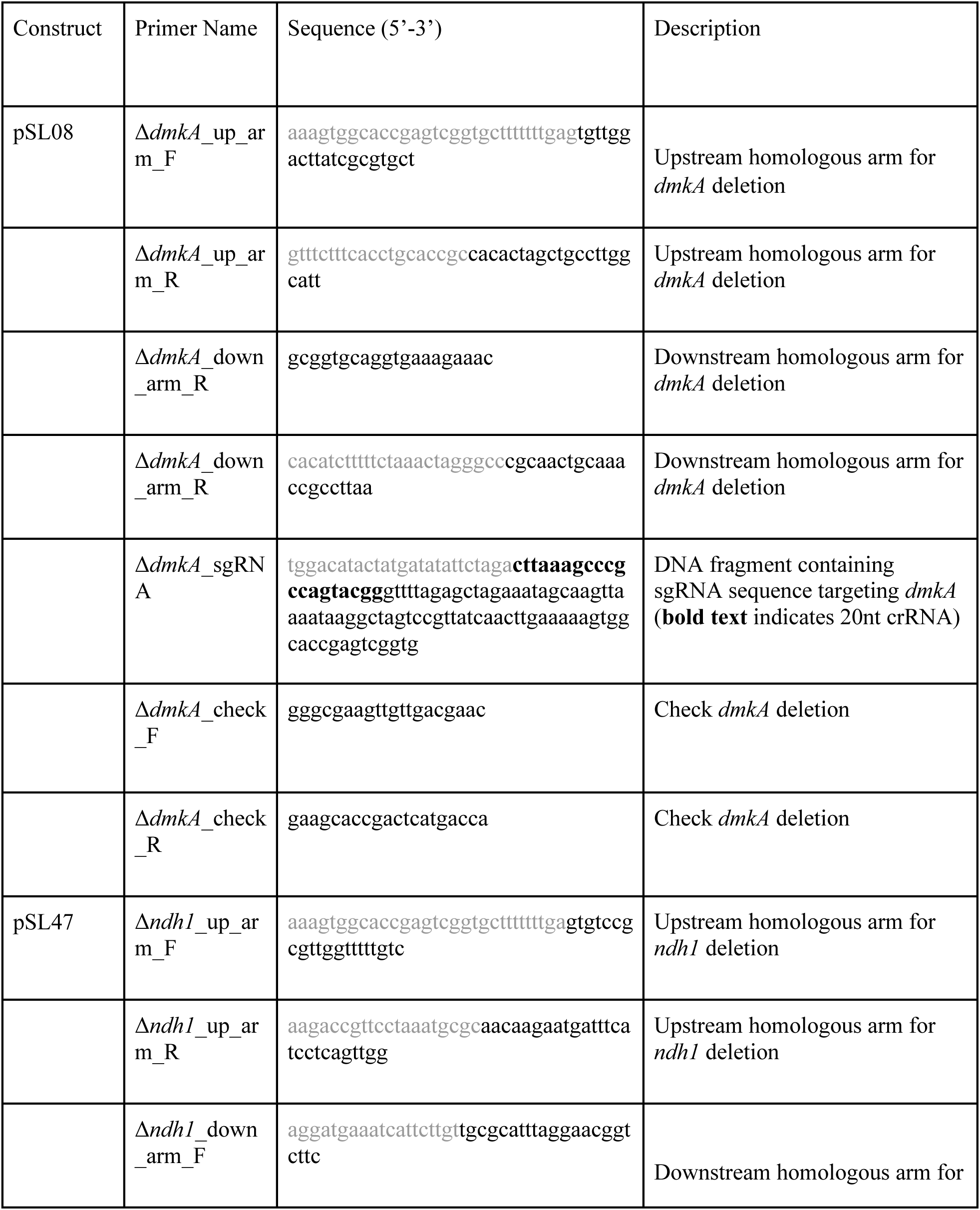
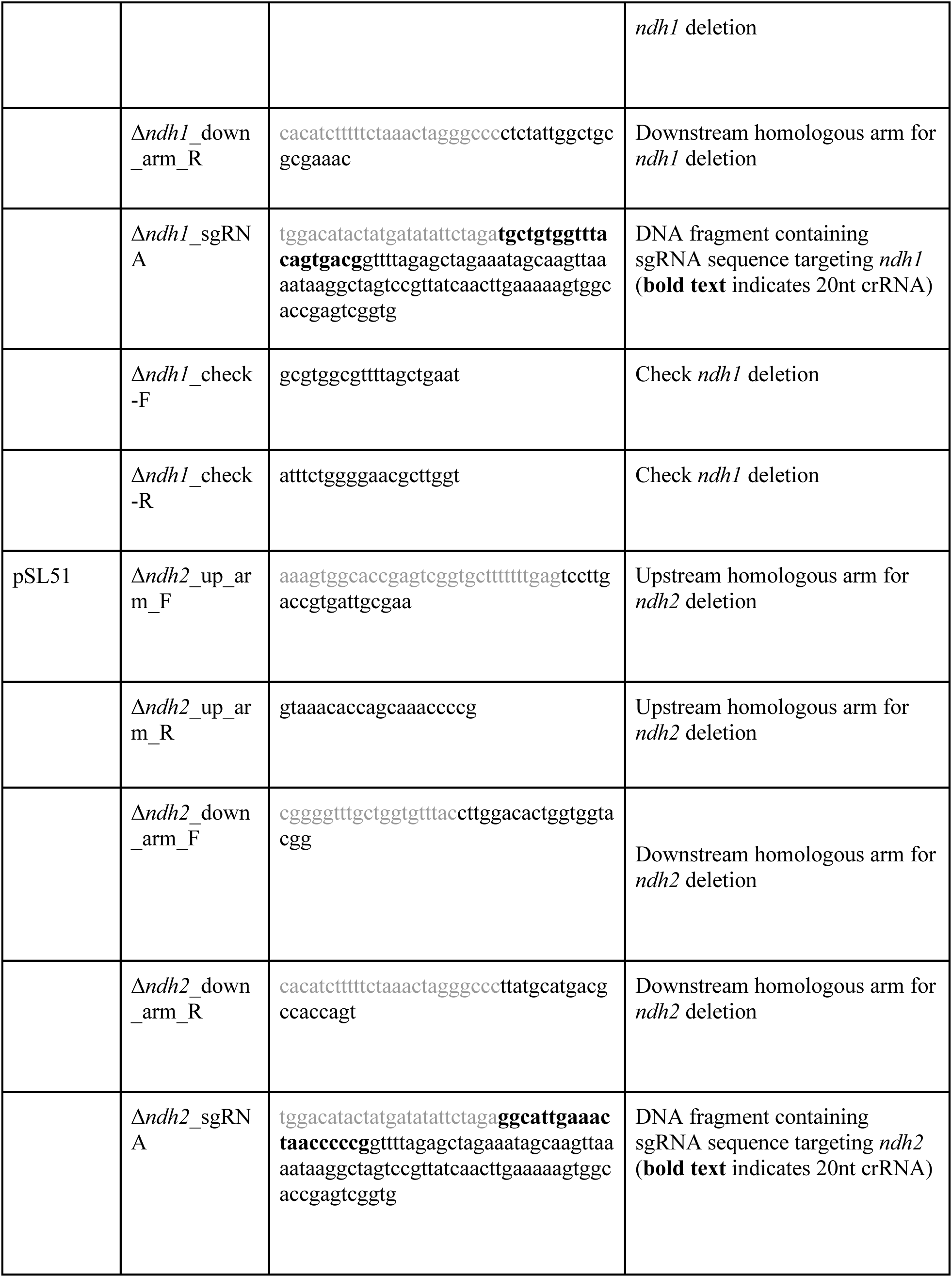
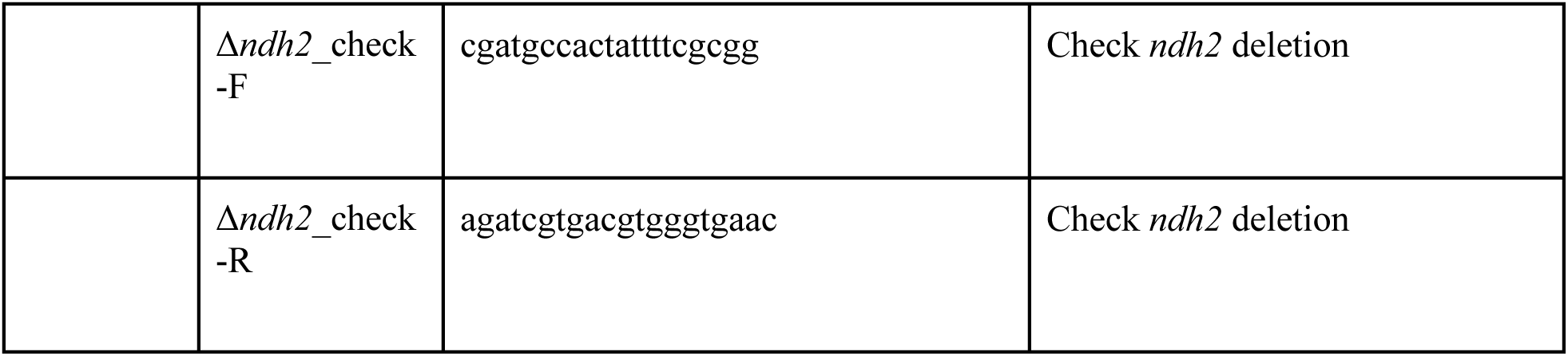
Primer used in this study.

